# labelSeg: segment annotation for tumor copy number alteration profiles

**DOI:** 10.1101/2023.05.17.541097

**Authors:** Hangjia Zhao, Michael Baudis

## Abstract

Somatic copy number alterations (SCNA) are a predominant type of oncogenomic alterations that affect a large proportion of the genome in the majority of cancer samples. Current technologies allow high-throughput measurement of such copy number aberrations, generating results consisting of frequently large sets of SCNA segments. However, the automated annotation and integration of such data are particularly challenging because the measured signals reflect biased, relative copy number ratios. In this study, we introduce *labelSeg*, an algorithm designed for rapid and accurate annotation of CNA segments, with the aim of enhancing the interpretation of tumor SCNA profiles. Leveraging density-based clustering and exploiting the length-amplitude relationships of SCNA, our algorithm proficiently identifies distinct relative copy number states from individual segment profiles. Its compatibility with most CNA measurement platforms makes it suitable for large-scale integrative data analysis. We confirmed its performance on both simulated and sample-derived data from The Cancer Genome Atlas (TCGA) reference dataset, and we demonstrated its utility in integrating heterogeneous segment profiles from different data sources and measurement platforms. Our comparative and integrative analysis revealed common SCNA patterns in cancer and protein-coding genes with a strong correlation between SCNA and mRNA expression, promoting the investigation into the role of SCNA in cancer development.

## 2 Introduction

Genomic instability is a nearly ubiquitous hallmark of cancer. Cancer cells often lose the ability to maintain genome integrity, but the molecular basis of genomic instability is not always clear [1]. One consequence of genomic instability is the occurrence of somatic copy number alterations (SCNA), which are changes in the copy number of chromosome segments from the regional allele count in somatic (i.e. post germline) tissues. SCNAs represent the by extent largest contributions to genomic variation in cancer, with genetic components affected by SCNA frequently conferring selective advantages to affected cells, thereby promoting cancer initiation and progression [2].

Various methods are employed to detect SCNAs, ranging from (molecular-)cytogenetic and locus-specific techniques such as karyotype analysis, interphase fluorescence in-situ hybridization (FISH), and spectral karyotyping (SKY) to genomic microarrays and next-generation sequencing (NGS) methods. However, achieving a comprehensive capture of all CNA information remains challenging with any individual approach, given the distinct detection biases and limitations inherent in different technologies and platforms. These differences manifest in various aspects, such as the upper and lower detection sensitivity for CNA events of differing sizes, the requirement for matched reference samples, and the capability to detect allele-specific CNA events [3], and are compounded by varying processing pipelines.

In the meta-analysis of large and heterogeneous SCNA datasets, a major challenge lies in interpreting and comparing segmented copy number profiles derived from raw intensity data obtained using techniques such as microarrays and NGS. Frequently, such segmented CNA profiles constitute the solely accessible data due to privacy concerns related e.g. to the exposure of single nucleotide polymorphism (SNP) data. Typically, available CNA data is represented through genomic segments with the relative abundance of the DNA expressed as the log R ratio (logR), calculated by taking the log2 of the ratio between the observed intensity of a sample and a reference intensity. Notably, it is a normalized metric, spotlighting changes in relative copy numbers rather than an absolute copy number at a given genomic location. However, such a representation leads to challenges when comparing SCNA profiles across datasets, which are further compounded by the fact that signal scale and noise levels can vary widely across samples due to variations in clonal sample purity, ploidy, experimental steps in bio-sample preparation, measurement platform, and other factors.

To overcome the challenges of inter-sample comparisons, some tools such as CNVkit [4] and VarScan2 [5] adopt fixed, empirical thresholds to classify segments with signals beyond these thresholds into “duplication” or “deletion” CNA categories. However, the selection of these thresholds can significantly impact the results of such analyses. Lower cut-off values improve the sensitivity of variant detection especially for samples with admixed non-cancer tissue but can lead to many false positive calls. In contrast, larger cut-off values may enhance the calling precision but risk overlooking true variants, thereby introducing systematic bias particularly due to cancer-specific differences in sample purity. Some studies [6, 7] optimize this process by incorporating purity estimation and adjusting thresholds based on the estimated purity. However, this added estimation complicates SCNA profiling by necessitating manual selection of the most suitable solutions [8]. Several complicated models have been developed to improve SCNA detection, including Gaussian mixture models [9, 10] and hidden Markov models [11, 12]. Although theoretically promising for providing more accurate CNA calls, these models often require raw data from individual measurement platforms that may not be accessible, or allele-specific data which certain technologies lack. Furthermore, most of these methods are designed to provide an absolute copy number quantification, rather than addressing the identification of distinct empirical CNA types. These types, characterized not only by CNA amplitude but also by CNA length, include broad CNAs and focal CNAs. These types differ in size, magnitude, and potential functional implications [13]. While GISTIC [14] can distinguish between these CNA types or levels, its primary objective is to identify recurrent amplified or deleted genomic regions across a set of samples, rather than accurately determining CNA levels in individual samples. Thus, the fast and accurate annotation of individual CNA segment profiles remains a challenging problem in the field of SCNA profiling. To address the above challenges, we developed a novel method called *labelSeg*. This method not only accurately annotates individual CNA profiles but also scales seamlessly to accommodate large-scale studies encompassing numerous samples. By utilizing estimated calling thresholds from individual segment profiles, *labelSeg* identifies various levels of CNAs without requiring prior information such as purity estimation. Its one-dimensional clustering approach, coupled with a direct cut-off strategy using the estimated thresholds, enables rapid processing of CNA profiles. The only input required is copy number segment profiles, which can be generated by most CNA measurement platforms and processing pipelines. These attributes make *labelSeg* particularly well-suited for large-scale meta-analyses. In validation cohorts from TCGA [15] covering diverse cancer types, *labelSeg* showed superior performance compared to GISTIC and fixed thresholds. Moreover, through an integrative analysis spanning 4 research projects, involving *>* 2000 glioblastoma samples and *>* 1200 lung squamous cell carcinoma samples, *labelSeg* demonstrated its capability to achieve fast, accurate, and comprehensive CNA profiling across a diverse and extensive collection of cancer samples. This achievement serves as a cornerstone for prospective comparative CNA analysis in cancer research.

### 3 Materials and Methods

*labelSeg* combines segment length and logR values to estimate appropriate thresholds for calling different levels of SCNA (Figure 1A). The comprehensive implementation of this algorithm is detailed in Figure 1B. There are several assumptions. First, the majority of detected copy number events are driven by predominant clones. It is a prevalent biological assumption that serves as the foundation for most contemporary algorithms utilized in the detection of SCNAs [16]. Under this assumption, segments of individual profiles form distinct clusters in logR values. These clusters are likely to represent different copy number states. Second, arm-level SCNAs are generally low copy number changes, whereas focal SCNAs can be of very high amplitude. This length-amplitude relationship of SCNA, which has been previously reported [13], allows reliable discrimination of different SCNA levels, including low-level duplication/deletion and high-level duplication/deletion.

**Figure 1:**
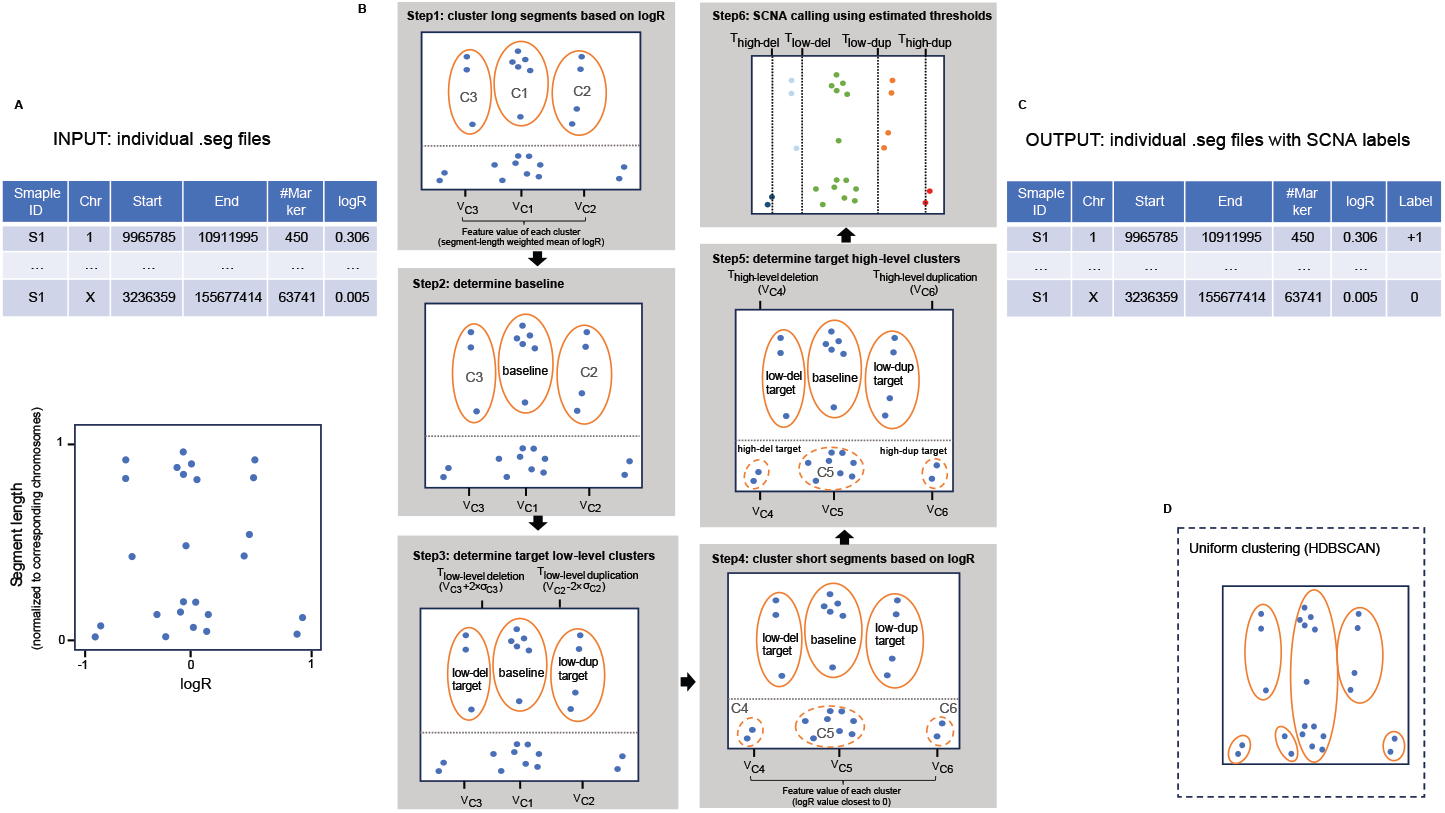
Workflow of *labelSeg*. (A) Input: The segmented data file provides segment lengths and associated numeric values (logR) for each segment, which are used in subsequent calculations. In the illustration below, each dot represents a single segment, with its normalized segment length displayed on the Y-axis and its logR value displayed on the X-axis (B) Algorithm: Step 1: Segments are categorized into long and short segments using a cutoff value (0.2) marked by a dashed horizontal line. Long segments are clustered in the logR space (X-axis) using DBSCAN, and these clusters are ranked based on the sum of segment lengths within each cluster. A feature value is computed for each cluster. Step 2: The baseline is determined using the feature value and rank. Step 3: Low-level target clusters are then identified using the distance of the feature value to the baseline and their respective ranks. Step 4: Short segments are also clustered in the logR space using DBSCAN with a larger radius, and these clusters are ranked based on corresponding feature values. Step 5: Highlevel target clusters are identified considering the distance of the feature value to the baseline, the predefined low-level target clusters, and their respective ranks. Calling thresholds are subsequently calculated based on these target clusters. Step 6: The cutoff determined by these estimated thresholds is applied to the segments. (C) Output: The original segment data file is augmented with an additional column of SCNA labels, representing relative copy number states. (D) Uniform clustering: A specialized clustering strategy employed in labelSeg when the clustering method utilizes HDBSCAN, an extension of DBSCAN.

Currently, the definitions of CNA magnitudes vary across different studies [17–23]. In general, high-level CNAs involve substantial changes in the copy number of chromosomal segments, with high-level duplication indicating the presence of multiple copies of certain genomic regions, and high-level deletion indicating either the complete loss or substantial reduction of specific regions. In contrast, low-level CNAs involve smaller changes in copy number compared to their high-level counterparts. In this study, we proposed the transformation of absolute copy numbers to relative CNA levels, grounded in a consensus derived from the reviewed studies. Specifically, we define high-level duplication as three or more gains compared to the ploidy, low-level duplication as one to two gains, high-level deletion as the absence of any copies, and low-level deletion as partial loss.

### 3.1 Algorithm

#### 3.1.1 Clustering

The segments in a sample are divided into long and short segments based on their length relative to the corresponding chromosomes. The criterion for segment size separation was determined based on the empirical distribution of segments derived from a combined dataset across various tumor types (Supplementary Figure S1). Segments occupying ≥20% of a chromosome are considered as long segments, and segments occupying *<* 20% of a chromosome are considered as short segments. Long and short segments are subjected to distinct clustering processes based on their logR values. The rationale underlying this choice is rooted in the intrinsic characteristics of long and short segments. Typically, long segments represent broad CNAs and tend to be derived from more measurement markers. As a consequence, long segments exhibit much lower variance and scales in logR values compared to short segments.

We employ the Density-Based Spatial Clustering of Applications with Noise (DBSCAN) [24] algorithm for clustering. There are two important parameters in DBSCAN: *ϵ* and *minPts*. The parameter *minPts* is the minimum number of points required to form a dense region. By default, this value is set to 1 to allow the formation of clusters even with a single segment (e.g. focal amplification), though it can be customized by users. The parameter *ϵ* is the maximum distance between two points for one to be considered in the neighborhood of the other. In our algorithm, we determine *ϵ* using a self-adaptive method. Initially, these values are set at predetermined levels (0.05 and 0.1 for long and short segment clustering, respectively). Subsequently, *ϵ* is systematically reduced by 0.01 until the standard deviations of logR in all clusters fall below specified thresholds (0.05 and 0.1 for long and short segment clustering, respectively). As a result of this variance control, it’s possible for segments sharing the same relative copy number state to form multiple clusters in logR values. While this phenomenon might potentially lead to an underestimation of the genuine logR variance within the same copy number state, this underestimation doesn’t hinder the determination of suitable thresholds. This resilience is attributed to both a variance adjustment (see Supplementary Data Section 1.1) and the utilization of specific “target clusters” in our threshold estimation procedure. These “target clusters” are meticulously identified through ranking methodologies that ensure precise threshold determination (elaborated further in the following sections).

Furthermore, our algorithm accommodates alternative clustering methods: Ordering Points To Identify the Clustering Structure (OPTICS) [25] and Hierarchical Density-Based Spatial Clustering (HDBSCAN) [26]. Both methods are extensions of the DBSCAN framework, simplifying the parameterization by removing the need to choose an appropriate *ϵ* value. HDBSCAN is a hierarchical extension for varying *ϵ* values and automatically determining the optimal number of clusters. It requires only the minimum cluster size (*minPts*) as input. Given that clustering by segment length intends to adapt to clusters of varying density while HDBSCAN is able to overcome such limitation, uniform clustering across all segments via HDBSCAN is reasonable and becomes a viable choice within our algorithm (Figure 1D). OPTICS replaces *ϵ* with an upper limit for the neighborhood size (typically rather high and set to infinity in *labelSeg*), generating an ordered list of data points such that points that are spatially closest become neighbors in the ordering. Although OPTICS doesn’t require the *ϵ* parameter, it does require a threshold to identify clusters from the OPTICS ordering. In our implementation, the value of the splitting threshold is set to the same value as the *ϵ* parameter in DBSCAN, resulting in the actual output of OPTICS comparable to DBSCAN. It is noted that the effect of the *minPts* parameter in OPTICS and HDBSCAN is different from its role in DBSCAN, as it not only defines the minimal cluster size but also contributes as a “smoothing” factor that involves density estimates.

#### 3.1.2 Calculate feature value for each cluster

Following the clustering process, a feature value, denoted as 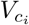, is computed for each segment group labeled as cluster *c*_*i*_. Depending on whether the cluster corresponds to long or short segments, the calculation methodology varies.

For clusters comprising long segments, the feature value is determined as the weighted mean of logR values from segments within the same cluster. Each of these logR values is assigned a weight based on its corresponding normalized segment length relative to the chromosome where it occupies. This weighted mean is used for calculating thresholds in identifying low-level SCNAs. In short segment clusters, the feature value is the logR value closest to 0 within the cluster, e.g. the minimum value in a cluster of positive logR values or the maximum value in a cluster of negative logR values. These feature values within the short segment clusters are instrumental in the derivation of thresholds for detecting high-level SCNAs.

#### 3.1.3 Estimate baseline

The baseline, representing segments with a neutral copy number state, is determined from the long segment clustering results. Specifically, the cluster that exhibits a feature value closest to 0 and occupies *≥* 40% of the total measured genomic region is considered as the baseline cluster, denoted as *c*_baseline_. In situations where data noise is pronounced or the profile is over-segmented, it is conceivable that no long segment clusters will span over 40% of the total length. In such instances, a step-wise reduction of the percentage limit, from an initial value of 40% to 20%, by a decrement of 10%, is performed until the baseline cluster is found.

During this step, users can interactively adjust the baseline upwards or downwards. The algorithm will subsequently identify the cluster that most closely aligns with the predefined baseline cluster while satisfying the aforementioned criteria.

#### 3.1.4 Estimate low-level calling threshold

The next step is to find clusters that represent low-level SCNA events. To increase tolerance to sub-clone effects, the remaining long segment clusters are ranked in descending order based on the cumulative segment length, denoted by {*c*_1_, *c*_2_, …, *c*_*k*_}. The target low-level clusters are determined as follows.

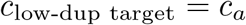

where,

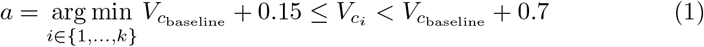

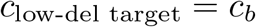

where,

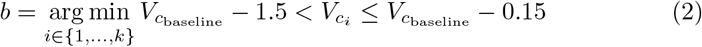

‘Target cluster’ is the cluster used to estimate calling thresholds. The lower bounds in Equations (1) and (2), set at ±0.15, aim to filter out noise from true SCNAs and can be tuned by users, although default values have been chosen based on empirical experience. The upper bounds are set to avoid calling high-level SCNAs and may also be adjusted based on prior knowledge. The default upper bound for duplication, 0.7, is derived from the theoretical logR value observed when the event of 5-copy gain occurs in 70% of diploid cells in the analyzed tissue. Similarly, the default upper bound for deletion, −1.5, is based on the theoretical logR value corresponding to a homozygous deletion event occurring in 70% of diploid cells in the measured tissue. When segment profiles exhibit high quality–usually indicative of high tumor purity and low measurement noise–employing higher bounds has the potential to improve the calling performance (Supplementary Figure S3B). To avoid overfitting, all results adhere to the default bounds.

To ensure calling sensitivity, the standard deviation of logR values from segments within the target cluster is included in the calculation of calling thresholds. Let T_low-level SCNA_ represent the calling threshold for low-level SCNA, and 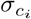 denote the standard deviation of logR in the cluster *c*_*i*_. The calculation proceeds as follows:

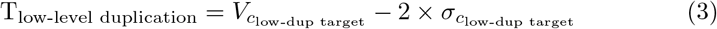

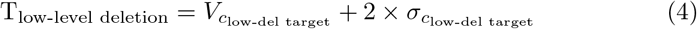

### 3.1.5 Estimate high-level calling threshold

The process of determining high-level calling thresholds integrates the clustering of short segments, the utilization of previously identified target low-level clusters, and the numeric correlation of logR distances among distinct copy number states. An interesting observation arises regarding the logR distances between low-level copy changes and neutral copy changes, as well as between high-level copy changes and neutral copy changes. These logR distances demonstrate a robust association with varying tumor sample purity and ploidy. Further insights into this phenomenon can be found in Supplementary Data Section 1.2 and Supplementary Figure S2. By leveraging this established correlation, as shown in Equations (5) and (6), the algorithm effectively finds a reliable calling threshold within heterogeneous samples to distinguish between high-level duplications/deletions and their low-level counterparts.

Short segment clusters are ranked by feature values in ascending order, denoted as {*c*_1_, *c*_2_, …, *c*_*m*_}. The target high-level clusters are determined as follows:

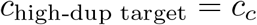

where,

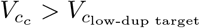

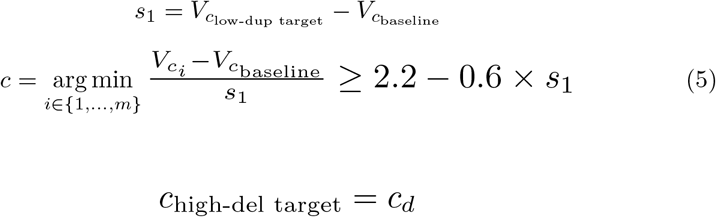

where,

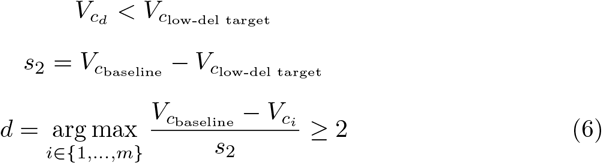

Given the unique characteristics of high-amplitude focal SCNAs and low-amplitude broad SCNAs, such as their different potential to pinpoint oncogenes and tumor-suppressor genes, the identification of high-level SCNAs is refined by calling only those focal SCNAs with amplitudes surpassing those of broad SCNAs. The calling threshold for high-level SCNAs is calculated as follows. Let **V**_**L**_ be the set of logR values of all previously defined long segments,

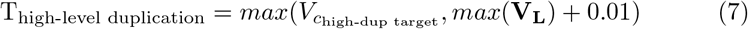

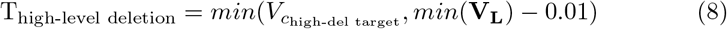

#### 3.1.6 SCNA Calling

Once the estimation of calling thresholds is complete, the subsequent step is to assign labels indicating relative copy number states or SCNA levels to each segment. The classification is carried out according to the following principles:

- Segments with logR ≤ T_high-level del_ are labeled as “-2”, indicating high-level deletion. If the sample is diploid, the high-level deletion is a homozygous deletion.
- Segments with logR *>* T_high-level del_ and ≤ T_low-level del_ are labeled as “-1”, meaning low-level deletion.
- Segments with logR ≥ T_low-level dup_ and *<* T_high-level dup_ are labeled as “+1”, meaning low-level duplication.
- Segments with logR ≥ T_high-level dup_ are labeled as “+2”, meaning high-level duplication.
- Segments with logR *>* T_low-level del_ and *<* T_low-level dup_ are labeled as “0”, meaning no changes in total non-allelic copy numbers.

There are other exceptions. For example, in cases where the segment profile is over-segmented and no long segments are present, or only high-level focal SCNAs occur within a sample, specific strategies for handling these exceptions are elaborated in Supplementary Data Section 1.3.

### 3.2 Implementation

*labelSeg* was written under R and the software is available at https://github.com/baudisgroup/labelSeg. The average execution time under default parameters was approximately 0.005 seconds per sample with 200 segments on a MacBook Pro18,4 (Apple M1 Max, 10 cores, 64 GB RAM) using 1 core.

## 4 Results

### 4.1 Performance on simulated data

The assessment of *labelSeg*’s performance involved a systematic examination of different clustering methods and variations in the *minPts* parameter, as depicted in Supplementary Figure S3A. The evaluation was conducted using simulated segment data that encompassed different scenarios, including variations in sample purities and noise levels (refer to Supplementary Data Section 2.1 for data generation details). The results from this benchmark indicated a limited impact of parameter choices on the calling performance. Consequently, our attention here is focused on two key configurations within *labelSeg* that were chosen to be representative: distinct DBSCAN clustering with *minPts* set to 1 for both short and long segments, and uniform HDBSCAN clustering with *minPts* set to 10. These selections were made based on their specific advantages. The former configuration offers computational simplicity, while the latter leverages the strengths of HDBSCAN, resulting in a specialized clustering strategy (illustrated in Figure 1D). Importantly, these variations do not affect other crucial steps in the algorithm, including individual feature value computation, cluster ranking, and threshold determination for both low and high-level SCNAs.

In the simulated scenario with moderate noise, we compared the performance of *labelSeg* with the application of optimal thresholds tailored to specific purity levels (Figure 2). The results were consistent with expectations: optimal thresholds demonstrated superior overall performance when matched with the true tumor purity they were designed for. However, the effectiveness of these thresholds diminished when applied to samples deviating substantially from the target purity. Conversely, *labelSeg* consistently exhibited commendable F1 scores across all simulated scenarios, showcasing its robustness across a broad spectrum of tumor sample purity levels. Notably, the two parameter sets within *labelSeg* showed subtle differences. DBSCAN clustering with *minPts* set at 1 exhibited enhanced adaptability in scenarios characterized by low sample purity, whereas HDBSCAN uniform clustering with *minPts* set at 10 excelled in situations of higher sample purity. Even in scenarios with exceptionally high noise levels, all compared methods displayed compromised performance, but this conclusion remained unchanged (Supplementary Figure S3A).

**Figure 2:**
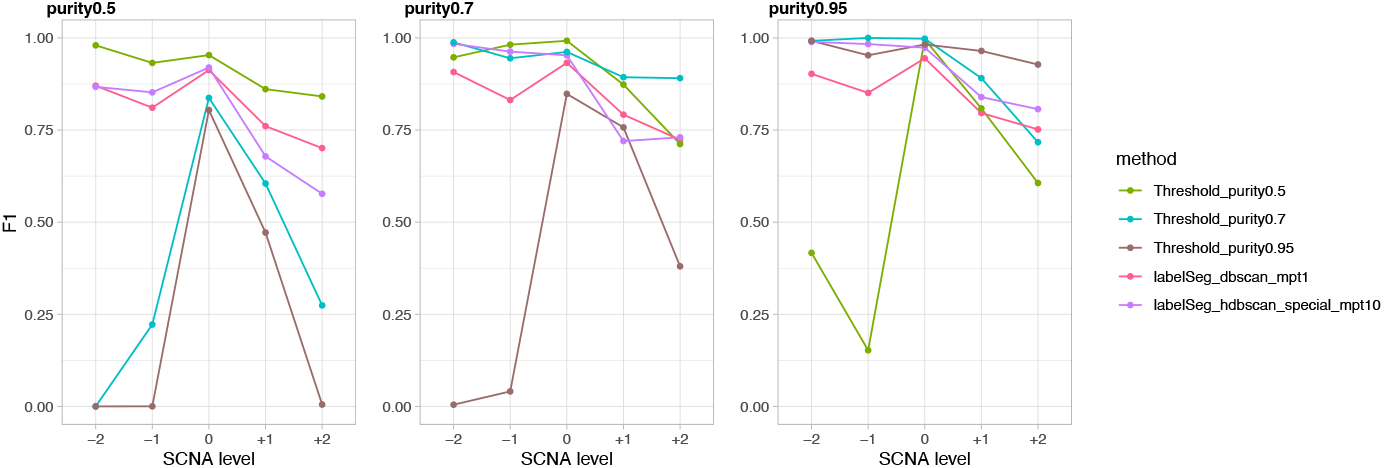
Performance on simulated data. The average F1 scores across samples were calculated for different levels of SCNA calling.

### 4.2 Performance on real data

To evaluate the performance of *labelSeg* on real biological data, we analyzed two TCGA datasets comprising 596 glioblastoma (GBM) and 503 lung squamous cell carcinoma (LUSC) samples. This evaluation involved a comparison with alternative methodologies, including GISTIC, a set of more stringent thresholds {±0.3, ±1 }, and a set of more relaxed thresholds {±0.15, ±0.7} (Figure 3). It’s worth noting that due to GISTIC’s gene-level profiling approach for relative copy number states, we conducted a conversion from segment-level labels to genelevel labels for both *labelSeg* outputs and fixed threshold callings. The rationale for excluding GISTIC from simulated data validation stems from GISTIC’s consideration of functional genomic elements in the genome and their varying background rates of SCNAs, a facet not accounted for in the simulation. Consequently, GISTIC’s applicability was limited in the simulation context. Given the inherent challenge in determining absolute truth within real biological data, we used ASCAT [27] estimates as a benchmark reference. These estimates provide gene-level absolute copy numbers extracted from the same datasets, which are then converted into relative CNA levels per gene. While ASCAT may not be a gold standard, it benefits from leveraging allelic information, a dimension not included in the benchmarked methods. As a result, the alignment with ASCAT calls offers a glimpse into the proficiency of SCNA amplitude profiling. Further details regarding the conversion of SCNA label from segment to gene can be found in Supplementary Data Section 2.2.

**Figure 3:**
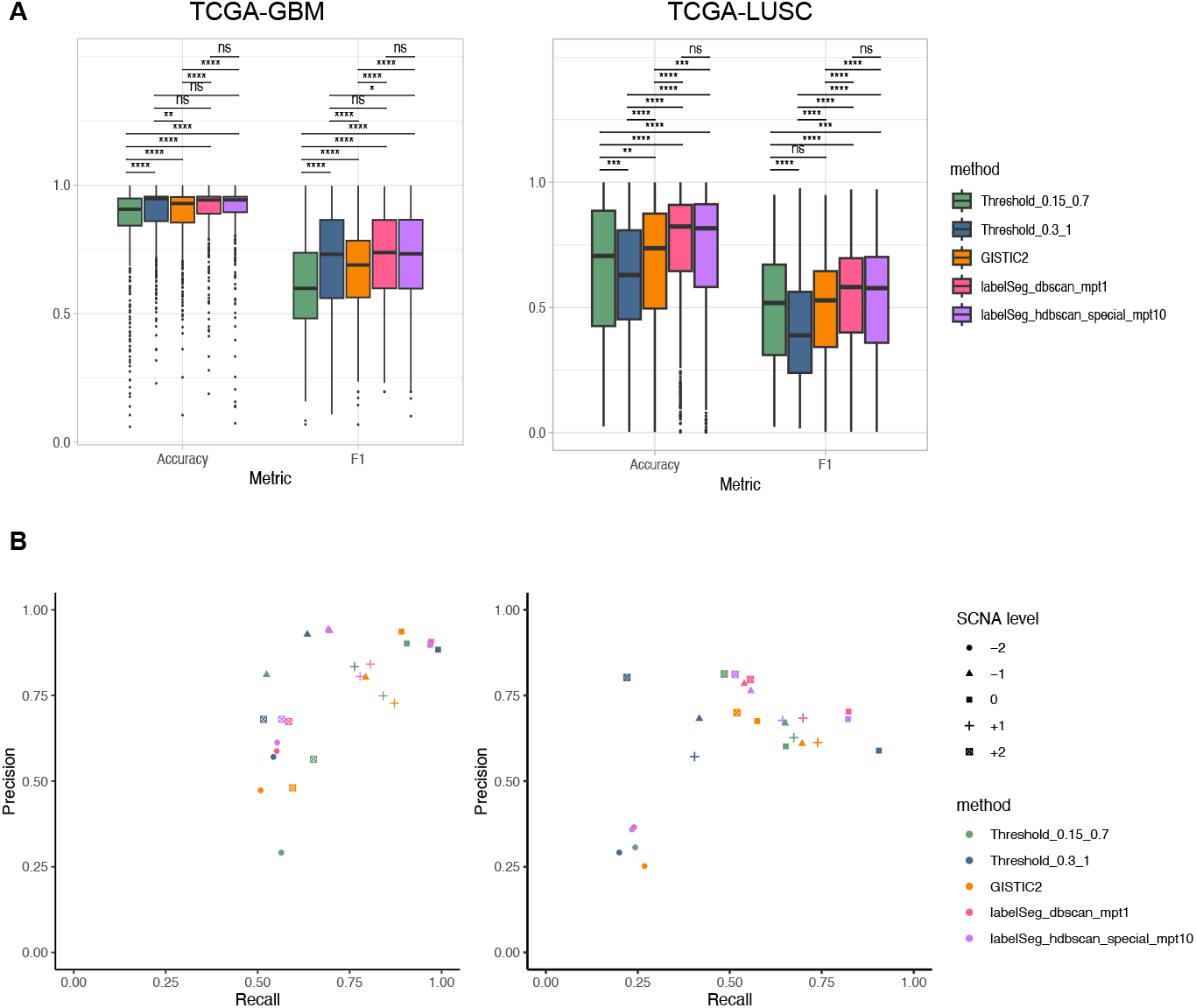
Performance on TCGA datasets. (A) Macro average of F1 scores across different SCNA classes and overall accuracy for each sample. (B) Average precision and recall across samples when calling different levels of SCNA.

In the GBM cohort, *labelSeg* and the stringent threshold (Threshold 0.3 1) had superior performance in terms of F1 score and accuracy (paired Wilcoxon rank sum test) (Figure 3A). In contrast, the relaxed threshold (Threshold 0.15 0.7) had the worst performance, primarily due to compromised precision (Figure 3B). Transitioning to the LUSC cohort, *labelSeg* once again demonstrated its prominence in terms of F1 score and accuracy, followed by GISTIC and the relaxed threshold. Interestingly, the stringent threshold exhibited limitations in this dataset due to issues of recall, unlike its performance in the GBM dataset. This underscores the inherent challenge posed by fixed thresholds in accommodating diverse tumor purities. As indicated by previous research, the average sample purity in GBM samples tends to surpass that in LUSC samples [28], which may underlie the performance discrepancy across the two datasets observed in all benchmarked methods. Remarkably, *labelSeg* revealed its strengths by outperforming all alternative methods in handling the more intricate dataset. Moreover, *labelSeg* showed similar performance across the two representative parameter sets (additional benchmarking results can be found in Supplementary Figure S4), confirming the negligible impact of parameterization. Considering factors such as algorithm complexity and robustness to complicated samples, we have designated separate DBSCAN clustering for both short and long segments with a *minPts* value of 1 as the default parameters for this method.

### 4.3 Integrative analysis of heterogeneous SCNA profiles

By design, *labelSeg* could provide a reliable calling no matter how heterogeneous the data are in terms of measurement platforms (input is segment file), tumor sample purities, and noise (clustering-based estimation). To illustrate its potential for precise and robust SCNA profiling—particularly in discerning between low-level broad SCNAs and high-level focal SCNAs—we applied this method to datasets originating from different resources and performed an integrative analysis. All of the ensuing results are generated using default parameters. There are four data resources from which the data was derived: TCGA, Progenetix [29], Clinical Proteomic Tumor Analysis Consortium (CPTAC) [30], and Cancer Cell Line Encyclopedia (CCLE) [31]. These datasets were generated through various measurement platforms and sample types, with detailed information provided in Table 1 and Supplementary Data Section 3.

**Table 1:** Summary of the glioblastoma segment datasets.

| Dataset | Sample | Sample type | Platform |
| --- | --- | --- | --- |
| TCGA | 596 | patient tumor samples | Affymetrix<br>SNP 6.0 |
| Progenetix | 1390 | patient tumor samples<br>and cancer cell lines | CGH array and<br>SNP array |
| CPTAC | 97 | patient tumor samples | WGS |
| CCLE | 64 | cancer cell lines | WES and WGS |

In glioblastoma samples, the SCNA patterns from different datasets were relatively consistent (Figure 4). Low-level SCNA frequency was calculated from segments with labels “+1” and “-1”. Chromosome 7 duplication and chromosome 10 deletion were the most prominent low-level SCNA features in glioblastoma samples with occurrence greater than 50% in all datasets and reaching 75% in the TCGA and CPTAC datasets. Chromosomes 9p, 13, 14 deletions and chromosomes 19, 20 duplications occurred in more than 25% of samples in most datasets. The UMAP plots based on the called low-level SCNA coverage of individual samples (Supplementary Figure S5) reveal the association of SCNAs across the genome. The co-occurrence of chromosome 7 duplication and chromosome 10 deletion was frequent in the analyzed samples. Duplications of chromosomes 19 and 20, and deletions of chromosome 9p were more likely to occur in samples with simultaneous chromosome 7 duplication and chromosome 10 deletion (*P*-value *<* 3e-12 for chr19, *P*-value *<* 9e-06 for chr20, *P*-value *<* 0.003 for chr9p, Pearson’s Chi-squared test). The SCNA pattern observed in the CCLE dataset was characterized by greater heterogeneity and noise compared to other datasets. This could possibly be explained by the difference between tumor samples and cell line models [32].

**Figure 4:**
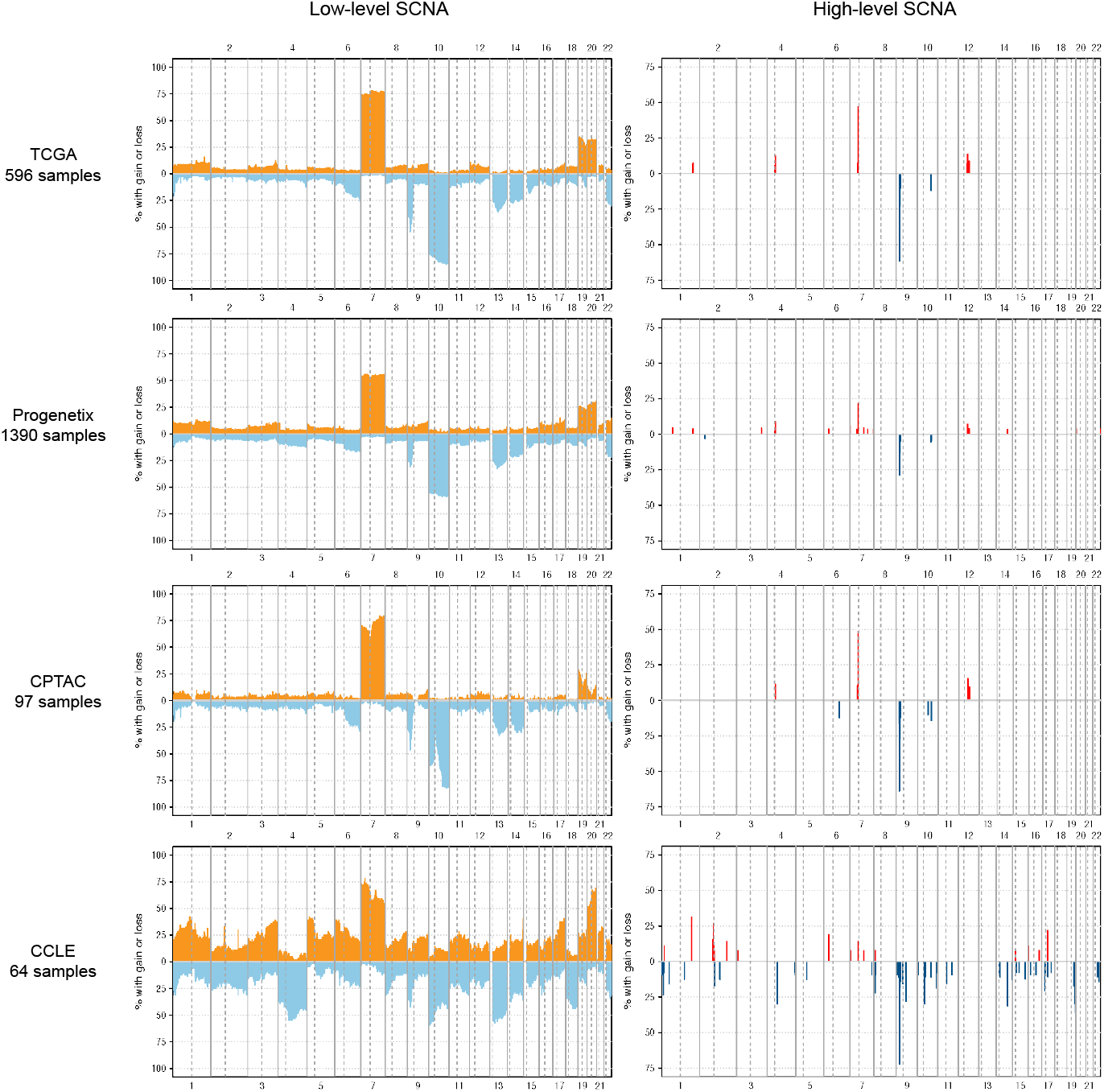
Frequency of SCNA calls in different glioblastoma datasets. Orange and red colors represent duplications, while light blue and dark blue colors represent deletions. The Y-axis displays the percentage of samples with SCNA overlapping with 1MB-sized genomic bins. The X-axis denotes chromosome numbers. Low-level SCNAs are identified by segments labeled “+1” and “-1”, while high-level SCNAs are identified by segments labeled “+2” and “-2”. In the frequency plots of high-level SCNAs, background noise peaks were filtered out.

High-level SCNA frequency was calculated from segments with labels “+2” and “-2”. Multiple consensus high-level SCNA focal peaks were observed across various analyzed projects, indicating the recurrent SCNAs’ robustness and reliability. Similar to low-level SCNA pattern, the CCLE samples were also more heterogeneous in high-level SCNAs. Supplementary Tables S1-S2 provide further information regarding congruent high-level duplication peaks from at least two projects and high-level deletion peaks from at least three projects. The most frequent amplification cross datasets happened in chr7: 54.1-56.1 MB with a frequency of around 46% in TCGA and CPTAC, 21% in Progenetix, and 12% in CCLE. The most frequent high-level deletion (probably homozygous deletion) occurred in chr9: 21-23 MB with a frequency around 62% in TCGA and CPTAC, 28% in Progenetix, and 70% in CCLE. The reduced high-level frequency detected in the Progenetix samples could potentially be attributed to the heterogeneity of the microarrays employed, which vary in their detection sensitivity of small CNAs, in conjunction with the diversity of sample types analyzed (Supplementary Figure S6). Amplification peaks were identified at chr6: 31.8-32.8 MB and chr7: 92.1-93.1 MB exclusively in the CCLE and Progenetix cohorts. Since all analyzed CCLE samples are cell lines, we hypothesized that amplification in these peaks is more likely to occur in glioblastoma cell lines. To test this hypothesis, we examined the composition of the Progenetix samples that exhibited such amplification. We found that a considerable proportion of the samples (37.6%) with the interesting amplification were cell lines, which is significantly higher than the overall proportion of cell lines in the Progenetix samples (6.3%). This observation was confirmed by a Pearson’s Chi-squared test with a *P*-value less than 2.2e-16, indicating a potential association between cell line samples and amplification in those loci.

We also analyzed the lung squamous cell carcinoma datasets from these data resources (Supplementary Figure S7). Low-level and high-level SCNA patterns in lung squamous cell carcinoma samples were roughly accordant across projects. Duplications of chromosomes 1q, 2p, 3q, 5p, 7, 8q and deletions of chromosomes 3p, 5q, 8p were characteristic low-level SCNA patterns in these samples with frequencies between 25% and 50%. Apart from focal peaks, high-level SCNAs spanned chromosomes 3q and 5p with amplification.

### 4.4 Relationship between copy-number dosage and mRNA expression

To further illustrate the practical utility of *labelSeg* and the utility of CNA levels, we examined the correlation between copy-number dosage and mRNA expression of protein-coding genes with recurrent high-level CNAs. Specifically, we analyzed paired mRNA expression data (normalized STAR read counts by TMM [33]) and CNA profiles of glioblastoma samples from the TCGA-GBM project. We only considered protein-coding genes that were amplified or highly deleted with a frequency of *>* 5% in TCGA-GBM samples, and that were located in the consensus focal regions mentioned in Section 4.3 and Supplementary Tables S1-S2.

Our results showed that 62 out of 68 frequently amplified genes had significantly increased mRNA expression when the SCNA level was “+2” compared to those with copy-neutral SCNAs labeled as “0” (Kruskal-Wallis test). Similarly, 21 out of 24 frequently high-level deleted genes had significantly decreased mRNA expression when the SCNA level was “-2” compared to those with SCNAs labeled as “0”. The most frequently high-level duplicated and high-level deleted genes in this cohort were EGFR and CDKN2A, respectively, and we observed a strong association between SCNA levels and mRNA expression for both genes (Figure 5A). Supplementary Figures S8-S9 provide box plots illustrating the correlation between SCNA and mRNA expression for other genes.

**Figure 5:**
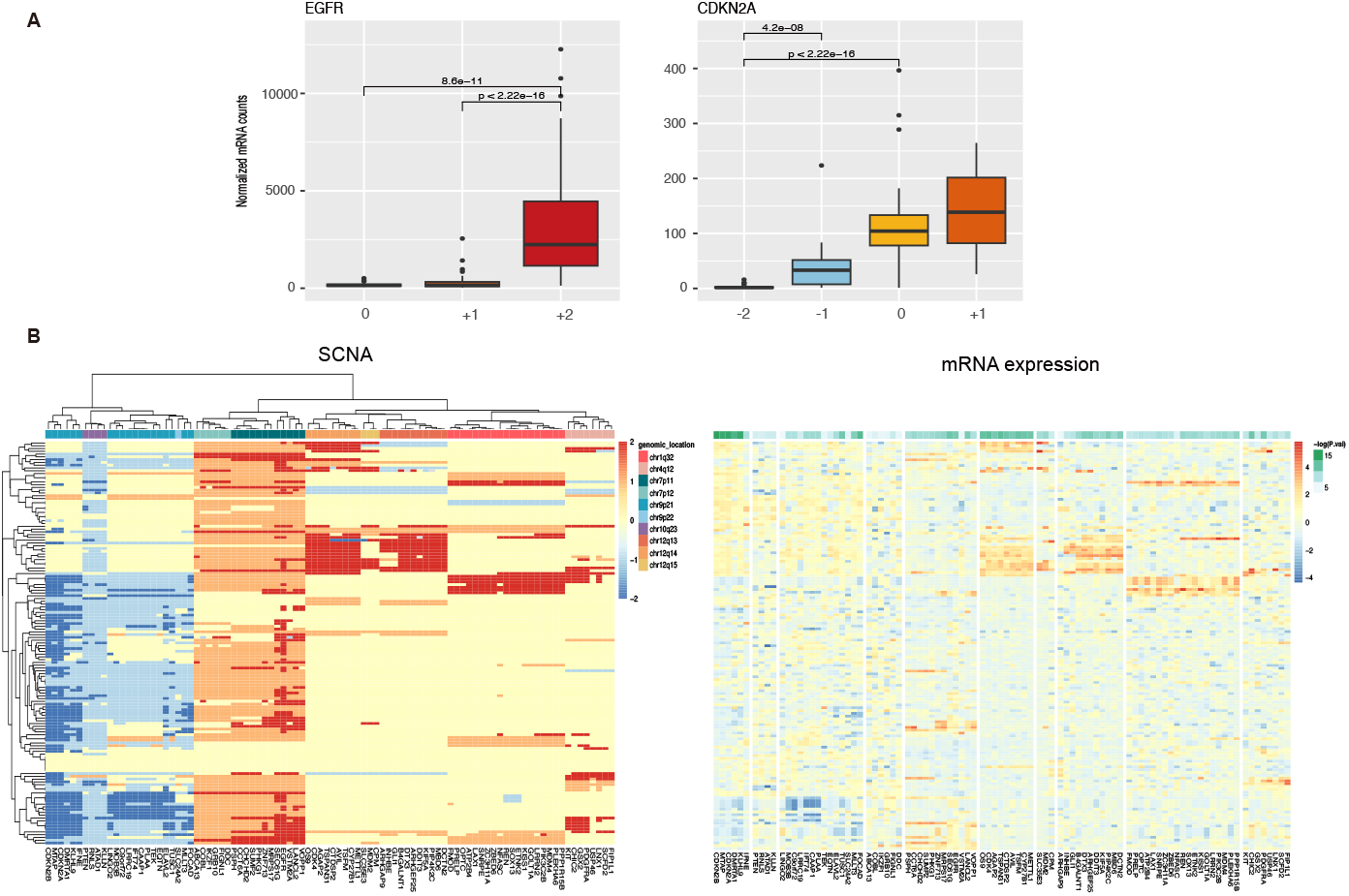
mRNA expression of genes with frequent high-level duplications and high-level deletions in glioblastoma. (A) mRNA expression of characteristic genes across different SCNA levels. (B) Heatmap displaying SCNA levels and mRNA expression in TCGA glioblastoma samples. Rows correspond to samples, and columns correspond to genes. The left SCNA heatmap presents SCNA label values. The right mRNA heatmap maintains the same row and column order as the left plot. P values were computed using the Kruskal-Wallis rank sum test and BH-adjusted from TMM-normalized mRNA counts. These normalized mRNA counts were log-transformed and standardized across samples per gene for visualization.

Figure 5B further shows the high correlation between SCNA levels and mRNA expression of the frequently altered genes in copy number dosage. These genes were clustered based on their genomic location in SCNA levels, which is not surprising since SCNA is a large-scale genomic variation affecting multiple genes. Genes in cytobands chr7p11, Chr9p21, chr12q13, and chr12q14 were strongly regulated in mRNA expression by the relative copy number states. Interestingly, VSTM2A and ARHGAP9 were not influenced by SCNA levels, although mRNA expression of nearby genes was strongly impacted by SCNA. Enrichment analysis was performed separately for these frequently high-level duplicated and deleted genes (Supplementary Figures S10). Both sets of genes were enriched in the glioma signaling pathway, indicating the important role of high-level SCNA in cancer development.

A similar analysis was conducted on the TCGA-LUSC cohort. Of the 1383 frequently amplified genes, 938 had an associated mRNA expression with their SCNA levels, while 20 of the 25 frequently high-level deleted genes had an associated mRNA expression with their SCNA levels. Notably, the high-level deleted genes with mRNA expression responsible for copy number changes were also located in cytoband chr9p21, which contains CDKN2A and CDKN2B (Supplementary Figures S11). Several frequently amplified genes that did not show mRNA association were enriched in epidermal keratinocyte differentiation (Supplementary Figures S12).

## 5 Discussion

Comprehensive profiling of somatic copy number alterations is valuable for advancing our understanding of cancer development and improving precision medicine applications. Here, we provide a novel method *labelSeg* for fast and accurate annotation of CNA segments. Our method estimates thresholds for calling different CNA levels from individual segment profiles, allowing more complete and robust identification of SCNAs in heterogeneous samples. The use of separate clustering by segment length not only adapts to the biological and technical variance in both focal and broad segments but also increases the tolerance to sub-clone effects and noise. Compared to the use of fixed cut-off values, *labelSeg* achieves a similar calling speed but increased accuracy. Furthermore, it does not require additional estimation such as tumor sample purity or other prior knowledge, making it a more convenient and powerful tool for large-scale comparative and integrative analysis to overcome bias from individual studies or platforms.

Our study demonstrated that *labelSeg* outperformed previous methods in SCNA profiling, as evidenced by its higher accuracy and F1 score. The robustness of *labelSeg* was demonstrated by the consistent patterns of SCNAs detected across heterogeneous datasets, making it a suitable tool for integrative analyses. Our analysis further confirmed simultaneous chromosome 7 duplication and chromosome 10 deletion in glioblastoma samples, which has been previously reported in other studies, thus highlighting the detection accuracy of genome-wide low-level CNAs. Furthermore, we identified several consensus high-level SCNA focal peaks enriched in protein-coding genes, which were observed in at least two of the four datasets. For most of these genes, mRNA expression strongly correlated with the called SCNA status, providing insights into the impact of SCNA of driver genes on tumor evolution.

Because *labelSeg* solely requires the logR values of segments as input, it is compatible with any technology that delivers segment data from its processing pipeline, regardless of the original data type (i.e. count or intensity-based). However, this general compatibility to a wide range of e.g. microarray and NGS platforms arrives with certain limitations of the method, particularly in estimating absolute copy number and ploidy which require some information about allelic composition. Although this is not a problem when relative copy number states are targeted, it renders the method unsuitable for some investigations that require accurate absolute quantification of copy numbers, such as assessing the degree of aneuploidy in tumor cells. Also, while sensitivity to probe-level noise affects all calling methods to various degrees, in *labelSeg*, such noise can diminish the clustering of segments in the logR values, which can hinder the ability to set accurate calling levels.

To conclude, our study presents a new strategy for segment classification and annotation, which enhances the interpretation of heterogeneous segment profiles with respect to calling efficiency, accuracy, and granularity. The stratification of SCNAs based on distinct levels highlights their varying sizes, amplitudes, and potential functional roles in the context of cancer pathogenesis. As the role of SCNA information expands within cancer genomics applications, including clinical diagnostics and tumor classification [34–36], our dedicated profiling method presents a valuable tool for investigating the intricate relationship between SCNAs and tumor diagnoses, oncogenomic subtypes, and clinical outcomes.

## Acknowledgements

We thank all members of the Baudis group for contributions to Progenetix resource. Some of the results published here are based upon data generated by the TCGA Research Network: https://www.cancer.gov/tcga.

## References

(1) Negrini, S.; Gorgoulis, V. G.; Halazonetis, T. D. Nature reviews. Molecular cell biology 2010, 11, 220–228.

(2) Mustjoki, S.; Young, N. S. New England Journal of Medicine 2021, 384, 2039–2052.

(3) Zarrei, M.; MacDonald, J. R.; Merico, D.; Scherer, S. W. Nature Reviews Genetics 2015, 16, 172–183.

(4) Talevich, E.; Shain, A. H.; Botton, T.; Bastian, B. C. PLOS Computational Biology 2016, 12, e1004873.

(5) Koboldt, D. C.; Zhang, Q.; Larson, D. E.; Shen, D.; McLellan, M. D.; Lin, L.; Miller, C. A.; Mardis, E. R.; Ding, L.; Wilson, R. K. Genome Research 2012, 22, 568.

(6) Zack, T. I. et al. Nature Genetics 2013 45:10 2013, 45, 1134–1140.

(7) Davoli, T.; Uno, H.; Wooten, E. C.; Elledge, S. J. Science 2017, 355, DOI: 10.1126/SCIENCE.AAF8399/SUPPL_FILE/AAF8399-DAVOLI-SM.PDF.

(8) Carter, S. L. et al. Nature Biotechnology 2012, 30, 413–421.

(9) Van De Wiel, M. A.; Kim, K. I.; Vosse, S. J.; Van Wieringen, W. N.; Wilting, S. M.; Ylstra, B. Bioinformatics 2007, 23, 892–894.

(10) Boeva, V.; Popova, T.; Bleakley, K.; Chiche, P.; Cappo, J.; Schleiermacher, G.; Janoueix-Lerosey, I.; Delattre, O.; Barillot, E. Bioinformatics 2012, 28, 423–425.

(11) Wang, K.; Li, M.; Hadley, D.; Liu, R.; Glessner, J.; Grant, S. F.; Hakonarson, H.; Bucan, M. Genome Research 2007, 17, 1665.

(12) Ha, G. et al. Genome Research 2014, 24, 1881.

(13) Beroukhim, R.; Mermel, C. H.; Porter, D.; Wei, G.; Raychaudhuri, S.; Donovan, J.; Barretina, J.; Boehm, J. S.; Dobson, J.; Urashima, M., et al. Nature 2010, 463, 899–905.

(14) Mermel, C. H.; Schumacher, S. E.; Hill, B.; Meyerson, M. L.; Beroukhim, R.; Getz, G. Genome biology 2011, 12, 1–14.

(15) Weinstein, J. N.; Collisson, E. A.; Mills, G. B.; Shaw, K. R.; Ozenberger, B. A.; Ellrott, K.; Shmulevich, I.; Sander, C.; Stuart, J. M. Nature genetics 2013, 45, 1113–1120.

(16) Tarabichi, M.; Salcedo, A.; Deshwar, A. G.; Ni Leathlobhair, M.; Wintersinger, J.; Wedge, D. C.; Van Loo, P.; Morris, Q. D.; Boutros, P. C. Nature methods 2021, 18, 144–155.

(17) Valsesia, A.; Rimoldi, D.; Martinet, D.; Ibberson, M.; Benaglio, P.; Quadroni, M.; Waridel, P.; Gaillard, M.; Pidoux, M.; Rapin, B., et al. PLoS One 2011, 6, e18369.

(18) Myllykangas, S.; Böhling, T.; Knuutila, S. In Seminars in cancer biology, 2007; Vol. 17, pp 42–55.

(19) Zhang, Y.; Chen, F.; Fonseca, N. A.; He, Y.; Fujita, M.; Nakagawa, H.; Zhang, Z.; Brazma, A., et al. Nature communications 2020, 11, 736.

(20) Waddell, N.; Pajic, M.; Patch, A.-M.; Chang, D. K.; Kassahn, K. S.; Bailey, P.; Johns, A. L.; Miller, D.; Nones, K.; Quek, K., et al. Nature 2015, 518, 495–501.

(21) Hogarty, M.; Brodeur, G. The genetic basis of human cancer 2001, 2, 115–28.

(22) Krijgsman, O.; Carvalho, B.; Meijer, G. A.; Steenbergen, R. D.; Ylstra, B. Biochimica et Biophysica Acta (BBA)-Molecular Cell Research 2014, 1843, 2698–2704.

(23) Jamal-Hanjani, M.; Wilson, G. A.; McGranahan, N.; Birkbak, N. J.; Watkins, T. B.; Veeriah, S.; Shafi, S.; Johnson, D. H.; Mitter, R.; Rosenthal, R., et al. New England Journal of Medicine 2017, 376, 2109–2121.

(24) Ester, M.; Kriegel, H.-P.; Sander, J.; Xu, X., et al. In kdd, 1996; Vol. 96, pp 226–231.

(25) Ankerst, M.; Breunig, M. M.; Kriegel, H.-P.; Sander, J. ACM Sigmod record 1999, 28, 49–60.

(26) Campello, R. J.; Moulavi, D.; Sander, J. In Pacific-Asia conference on knowledge discovery and data mining, 2013, pp 160–172.

(27) Van Loo, P.; Nordgard, S. H.; Lingjærde, O. C.; Russnes, H. G.; Rye, I. H.; Sun, W.; Weigman, V. J.; Marynen, P.; Zetterberg, A.; Naume, B., et al. Proceedings of the National Academy of Sciences 2010, 107, 16910–16915.

(28) Aran, D.; Sirota, M.; Butte, A. J. Nature communications 2015, 6, 1–12.

(29) Huang, Q.; Carrio-Cordo, P.; Gao, B.; Paloots, R.; Baudis, M. Database 2021, 2021.

(30) Ellis, M. J.; Gillette, M.; Carr, S. A.; Paulovich, A. G.; Smith, R. D.; Rodland, K. K.; Townsend, R. R.; Kinsinger, C.; Mesri, M.; Rodriguez, H., et al. Cancer discovery 2013, 3, 1108–1112.

(31) Nusinow, D. P.; Szpyt, J.; Ghandi, M.; Rose, C. M.; McDonald III, E. R.; Kalocsay, M.; Jané-Valbuena, J.; Gelfand, E.; Schweppe, D. K.; Jedrychowski, M., et al. Cell 2020, 180, 387–402.

(32) Domcke, S.; Sinha, R.; Levine, D. A.; Sander, C.; Schultz, N. Nature communications 2013, 4, 1–10.

(33) Robinson, M. D.; Oshlack, A. Genome biology 2010, 11, 1–9.

(34) Louis, D. N.; Perry, A.; Wesseling, P.; Brat, D. J.; Cree, I. A.; Figarella-Branger, D.; Hawkins, C.; Ng, H.; Pfister, S. M.; Reifenberger, G., et al. Neuro-oncology 2021, 23, 1231–1251.

(35) Zhang, N.; Wang, M.; Zhang, P.; Huang, T. Biochimica et Biophysica Acta (BBA)-General Subjects 2016, 1860, 2750–2755.

(36) Elsadek, S. F. A.; Makhlouf, M. A. A.; Aldeen, M. A. In Proceedings of the International Conference on Advanced Intelligent Systems and Informatics 2018 4, 2019, pp 198–207.

