## Supplementary material for "labelSeg: segment annotation for tumor copy number alteration profiles"

**Section1 Algorithm description**

**1.1 Standard deviation of low-level target clusters**

To enhance noise tolerance and the sensitivity for detecting true variants, the determination of low-level calling thresholds involves the utilization of the feature value $V_{c_{low-level target}}$ (segment-length weighted mean) and the standard deviation $\sigma_{c_{low-level target}}$ of the respective target cluster $c_{low-level target}$. It should be noted that due to a variance control implemented in the initial clustering step, the cluster standard deviation may potentially be underestimated. In certain cases involving segment profiles of subpar quality, the cluster standard deviation might be excessively large, leading to an impractical calling threshold calculation. Therefore, an adjustment for standard deviation becomes imperative.

In scenarios where the cluster standard deviation is either unavailable (owing to identical logR values within the target low-level cluster) or deemed too small ( $\sigma_{c_{low-level target}}< \frac{|V_{c_{low-level target}}-V_{c_{baseline}}|}{10}$ ) or excessively large ( $\sigma_{c_{low-level target}}> \frac{|V_{c_{low-level target}}-V_{c_{baseline}}|}{2}$), the value of $\sigma_{c_{low-level target}}$ will undergo an adjustment and be set to $\frac{|V_{c_{low-level target}}-V_{c_{baseline}}|}{10}$.

When employing HDBSCAN clustering across all segments, the adjustment of small standard variation is omitted due to the non-usage of variance control in this specific configuration.

**1.2 Relationship of logR values between different copy number states**

The logR value is influenced by the tumor sample's purity and ploidy, posing a challenge in accurately determining diverse relative copy number states within heterogeneous CNA profiles. To address this issue, *labelSeg* employs a numerical model that establishes relationships between logR values corresponding to distinct copy number, accommodating varying levels of tumor purity and ploidy. We validated this relationship through the following model, and the results of this validation are presented in Supplementary Figure S2. For a given segment i with copy number $CN_{i}$ from sample j, characterized by tumor cell ploidy $Ploidy_{j}$ and purity $Purity_{j}$, the corresponding logR value is computed as follows:

$$logR_{CN_{i}} = log_{2}\frac{Purity_{j} \times CN_{i} + \left( 1-Purity_{j} \right)\times2}{Purity_{j} \times Ploidy_{j} + \left( 1-Purity_{j} \right)\times2}$$

Eq.S 1

In this model, the baseline logR value is always 0, although this is generally not the case in real scenarios. To account for this difference, *labelSeg* incorporates the distance between low/high copy changes and the estimated baseline as a representation of the genuine logR value arising from corresponding copy changes in practical implementation.

**1.3 Special cases**

In certain scenarios, *labelSeg* operates differently to address specific challenges and constraints. If segment profiles are excessively fragmented and no individual segment covers more than 20% of a chromosome, *labelSeg* will issue a warning and terminate its operation. In such cases, users may merge segments prior to applying the algorithm.

In instances where the segment profiles are heavily influenced by noise, *labelSeg* may struggle to accurately identify the baseline cluster. Should this occur, the feature value of the baseline cluster is set to 0 and utilized in subsequent calculations.

When long segment clustering fails to identify any low-level target cluster, e.g. the absence of broad low-level SCNAs in the analyzed sample, *labelSeg* calculates low-level calling thresholds from short segment clustering using the same method but with a more lenient boundary.

In situations where both long and short segment clustering do not yield low-level target clusters, indicating the absence of both broad and focal low-level SCNAs, *labelSeg* employs fixed distances of 0.6 and 1 from candidate clusters to the baseline cluster to pinpoint target high-level duplication and deletion clusters, respectively.

**Section2 Validation**

2.1 **Synthetic datasets**

The generation of simulated data was based on Willenbrock and Fridlyand's work with a slight modification [1]. For each sample, ploidy was assumed to be 2, and the true copy number (CN) of each segment was drawn from a fixed state probability $\{0.05,0.15,0.55,0.15,0.05,0.05\}$ when CN$=\left\{ 0,1,2,3,4,5 \right\}.$ The lengths of segments with different copy number states were assigned by sampling from the empirical length distribution, which was established by binning logR values of the combined segment dataset (Supplementary Figure S1). The intervals were less than -1.2 (0 copies), between -1.2 and -0.3 (one copy), between -0.3 and 0.3 (two copies), between 0.3 and 0.7 (three copies), between 0.7 and 1.2 (four copies) and greater than 1.2 (five copies). The intervals were constructed to simulate the real segment length distribution rather than to indicate the real logR-copy number relationship. Unlike the original method, the length of segments with CN = 1,2,3 was scaled by 2.5,3,2.5 respectively. This modification accounts for the prevalence of normal segments or SCNAs with low copy number alterations spanning large contiguous genomic regions. Gaussian noise was applied with a variable standard deviation depending on the segment length. Let $P_{j}$ denote the proportion of tumor cells, namely sample purity, in one sample j, $L_{i}$ represent the length of a segment i, $CN_{i}$ be the copy number of a segment i, $L_{Q_{2}\left( CN_{i} \right)}$ denote the ${50}_{th}$ percentile of the length of segments with the same copy number as $CN_{i}$, and $L_{Q_{3}\left( CN_{i} \right)}$ denote the ${75}_{th}$ percentile of the length of segments with the same copy number as $CN_{i}$. The final logR value for segment i in sample j was computed as follows:

$$log2 \frac{P_{j}\times{CN}_{i}+\left( 1-P_{j} \right)\times2}{2} + N(0,\sigma_{i})$$

Eq.S 2

where,

$$\sigma_{i}=\left\{ \begin{aligned} 0.05 if L_{i}>L_{Q_{3}\left( CN_{i} \right)} \text{\& }CN_{i}\in\left\{ 1,2,3 \right\} \\ 0.075 if L_{Q_{2}\left( CN_{i} \right)}< L_{i}\leq L_{Q_{3}\left( CN_{i} \right)}\text{ \& }CN_{i}\in\left\{ 1,2,3 \right\} \# \\ 0.1 others \end{aligned} \right.$$

Eq.S 3

We simulated various scenarios, including distinct tumor sample purities (0.5, 0.7, and 0.95) and a highly noisy situation where $\sigma_{i}$ was three times the original value indicated in Eq. S3. Each scenario comprised 500 samples, each containing 200 segments. To compare with *labelSeg* outputs, true copy numbers were transformed to true relative copy number state labels.

We compared *labelSeg* with optimal thresholds across different sample purities. The optimal thresholds were as follows: for purity 0.5, {-0.8,-0.3,0.25,0.7}; for purity 0.7, {-1.5,-0.5,0.35,0.9}; and for purity 0.95, {-2,-0.8,0.4,1.1}.

2.2 **Real datasets**

The TCGA segment datasets were acquired using *TCGAbiolinks* R package [2]. These datasets consist of masked copy number segment profiles derived from primary tumors and generated through the Affymetrix SNP 6.0 platform. After randomly removing duplicate samples from the same patient, the TCGA-GBM dataset comprised 596 samples from patients with glioblastoma multiforme (GBM), while the TCGA-LUSC dataset contained 503 samples from patients with lung squamous cell carcinoma (LUSC). The gene-level ASCAT results were downloaded from the GDC data portal. The absolute copy numbers estimated by ASCAT were then transformed into relative copy number state labels. Ploidy was defined as the absolute copy number spanning most regions of the genome.

GISTIC outputs the same CNA state labels as *labelSeg* that indicate relative copy number states. We used available gene-level GISTIC calls of TCGA datasets from cBioportal database [3, 4] for comparison. Except for the existing gene-level calls from GISTIC and ASCAT, other methods, including *labelSeg*, produced labels for each segment. These segment-level labels were subsequently transformed into gene-level labels. The assignment of gene-level labels was based on the copy number state of the overlapping segment with the maximal severity, as proposed in a previous study [5].

**Section3 Data collection**

The collection of TCGA segment datasets was described in Section 2.2. The Progenetix segment datasets were collected from the arrayMap cohort of Progenetix database. Both TCGA and Progenetix data were generated from microarrays, and segments with fewer than 10 probes were excluded. For the CCLE segment datasets, we downloaded them from the depmap data portal 22Q4 release, and duplicate profiles from the same cell line model were excluded after manual quality check. The CPTAC segment datasets were collected from two resources. Glioblastoma segment profiles were downloaded from cBioportal, and lung squamous cell carcinoma segment profiles were downloaded from GDC data portal. The lung squamous cell carcinoma segment profiles were AscatNGS calls [6] with absolute copy numbers instead of logR values, and we transformed them into SCNA labels. The genome version of these data is hg38, except for the CPTAC glioblastoma data, which were converted from hg19 to hg38 using *segment_liftover* Python package [7]. The mRNA expression data of matched TCGA samples are obtained via *TCGAbiolinks* R package.

**Section4 Supplementary tables**

**Supplementary Table S1:** Amplification peaks across glioblastoma samples from multiple projects. The genomic location is determined based on genomic bins of 1MB size in the hg38 genome.

| Chromosome | location (MB) |
| --- | --- |
| 1 | 203.4-205.4 |
| 4 | 52-56 |
| 6 | 31.8-32.8 |
| 7 | 52.1-57.1 |
| 7 | 92.1-93.1 |
| 12 | 56.5-58.5 |
| 12 | 67.5-69.5 |

**Supplementary Table S2:** High-level deletion peaks across glioblastoma samples from multiple projects.

| Chromosome | location (MB) |
| --- | --- |
| 9 | 19-29 |
| 10 | 86.8-88.8 |

**Section5 Supplementary figures**

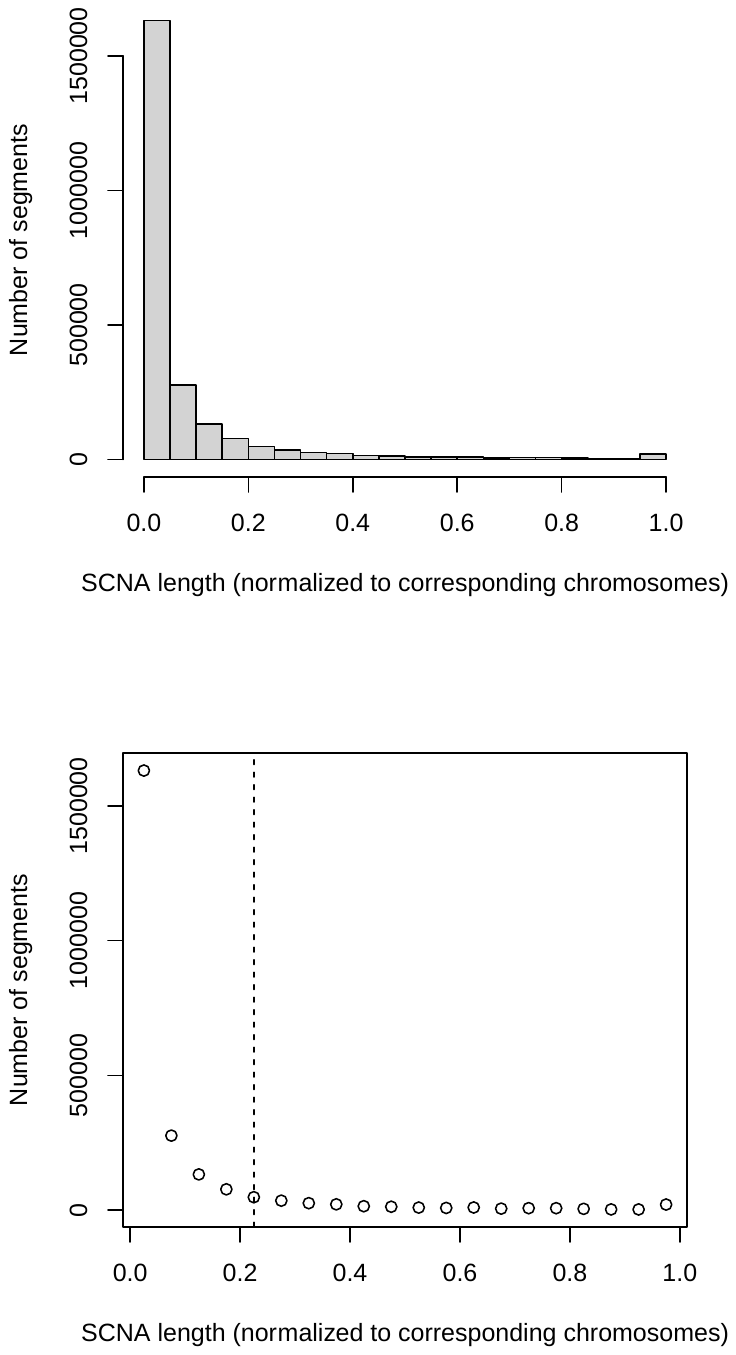

**Supplementary Figure S1: Length distribution of CNA segments across cancers.** The scatter plot depicts the data illustrated in the top histogram, with the elbow point indicated by a dashed line. The analyzed data comprise combined datasets of multiple cancer types from TCGA, including BLCA, BRCA, CESC, COAD, GBM, HNSC, KICH, KIRC, KIRP, LGG, LIHC, LUAD, LUSC, OV, SKCM, STAD, UCEC, and UCS.

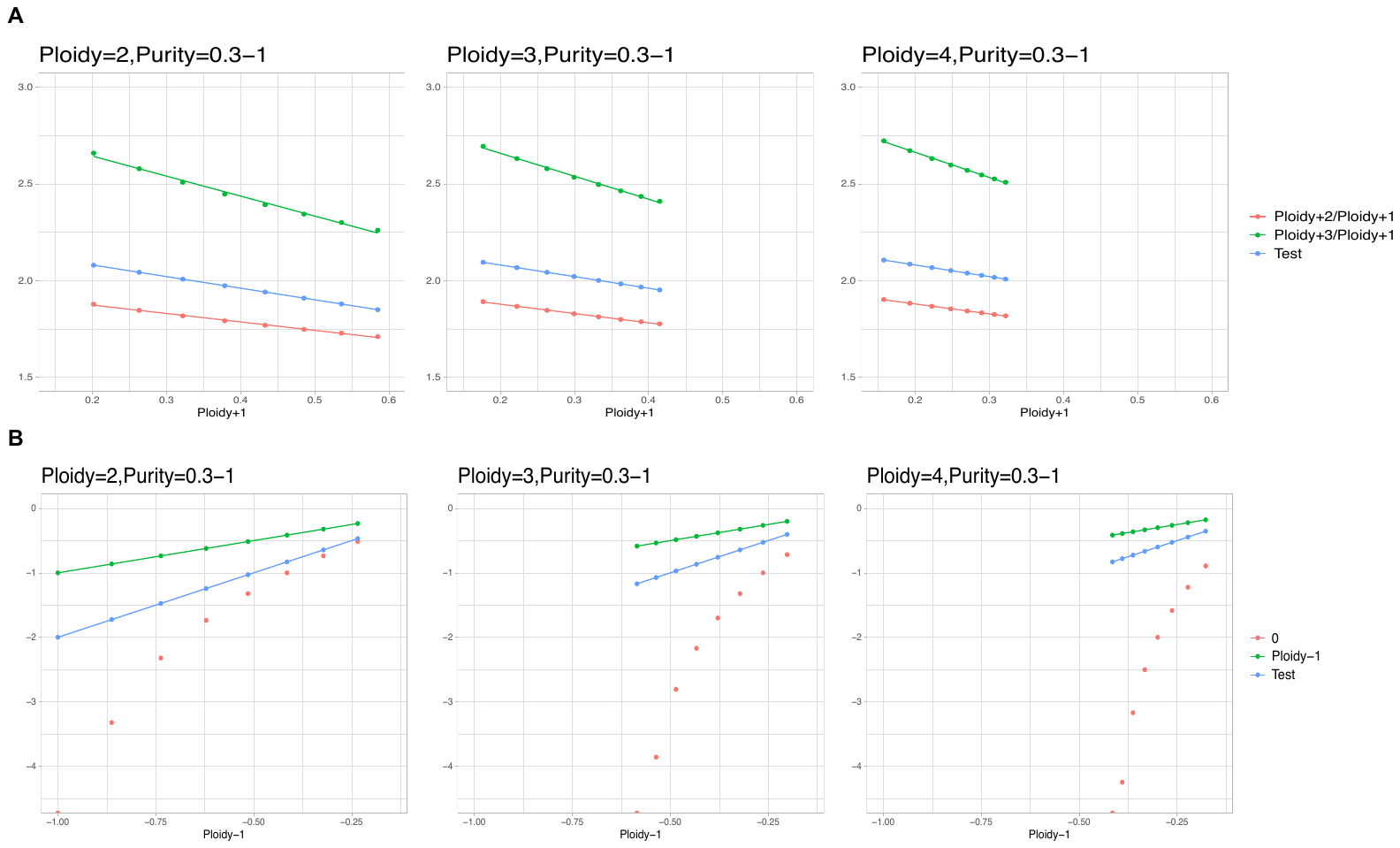

**Supplementary Figure S2: Validation of the logR relationship between different tumor copy numbers, ploidy, and purity.** logR calculation follows the model of Equation (S1)

(A) Duplication: The X-axis represents the logR value when the tumor cell copy number equals the tumor ploidy+1 (low-level duplication), under various tumor sample purities ranging from 0.3 to 1. The Y-axis indicates the ratio of logR values between different tumor cell copy numbers under the same sample purity as the X-axis. The "Test" value is -0.6x+2.2, which is utilized in determining high-level duplication calling thresholds, as explained in the main text Equation (5). The blue dots consistently fall between the red and green dots, indicating the ability to distinguish duplications with three or more copy gains (high-level duplication) from duplications with two or fewer copy gains (low-level duplication) relative to the overall ploidy.

(B) Deletion: The X-axis shows the logR value when the tumor cell copy number equals the tumor ploidy-1 (low-level deletion), under diverse tumor sample purities ranging from 0.3 to 1. The Y-axis represents the logR value at different tumor cell copy numbers under the same sample purity as the X-axis. The "Test" value is 2x, which is employed in finding high-level deletion calling thresholds, as demonstrated in the main text Equation (6). The blue dots consistently fall between the red and green dots, indicating the capacity to differentiate deletions with two or more copy losses (high-level deletion) from deletions with only one copy loss (low-level deletion) compared to the overall ploidy. Notably, when the ploidy is 2, *labelSeg* designates homozygous deletion as a high-level deletion.

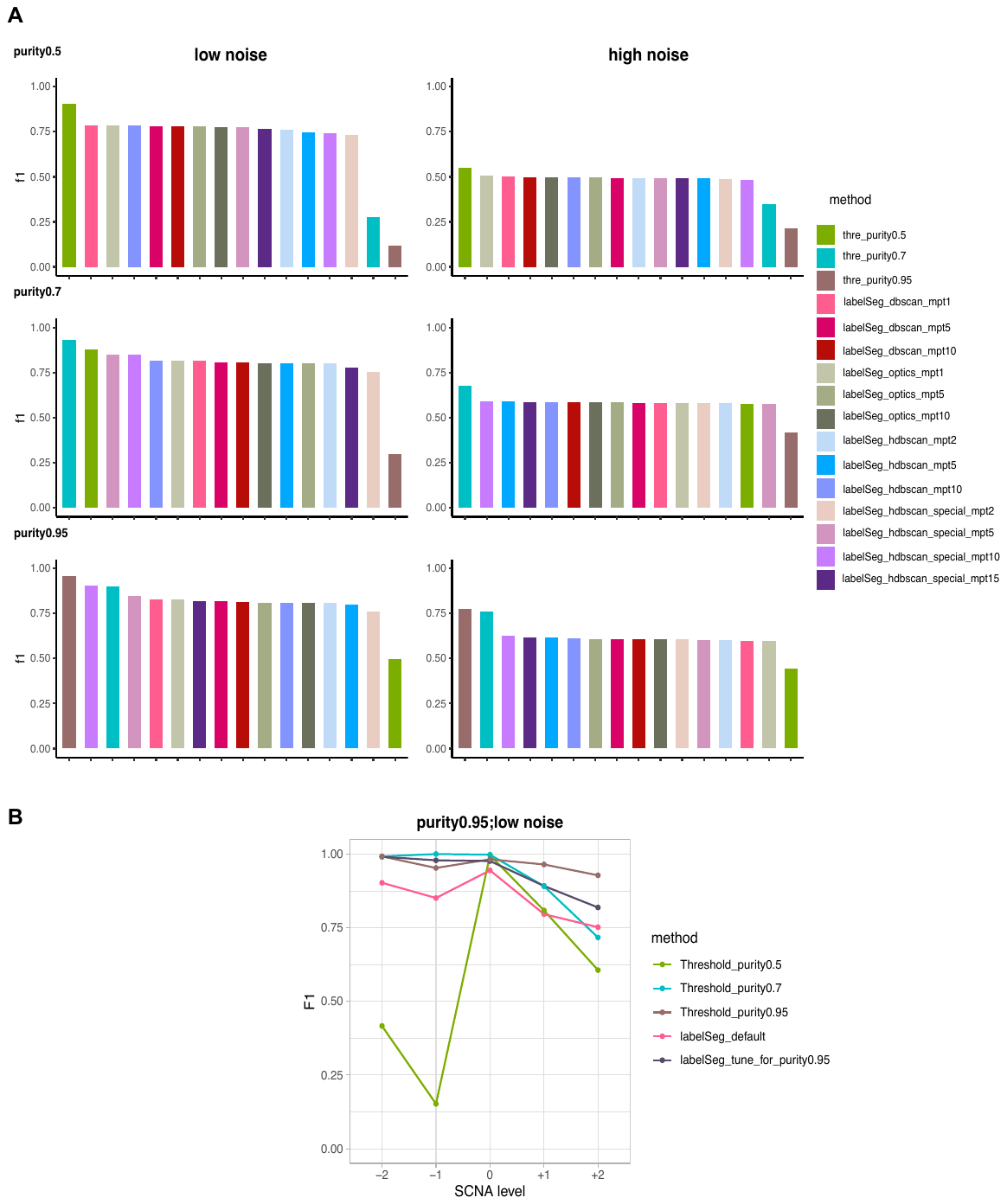

**Supplementary Figure S3: Benchmark performance on simulated data.** (A) Comparison between optimal thresholds for specific purity and *labelSeg* with tuning in clustering methods and minPts parameters. The Y-axis represents the average F1 scores across samples. For each sample, the F1 score was computed as the macro average of the F1 scores in calling low/high SCNA levels. (B) Comparison between optimal thresholds for specific purity and *labelSeg* with tuning in bounds. The Y-axis indicates the average F1 scores across samples. "labelSeg_default" utilized default bounds, where the lower bounds for duplication and deletion are 0.15 and -0.15; upper bounds for duplication and deletion are 0.7 and -1.5. "labelSeg_tune_for_purity0.95" used higher bounds than default, where the lower bounds for duplication and deletion are 0.3 and -0.6; upper bounds for duplication and deletion are 1 and -2. Other parameters were kept the same (DBSCAN, minPts1).

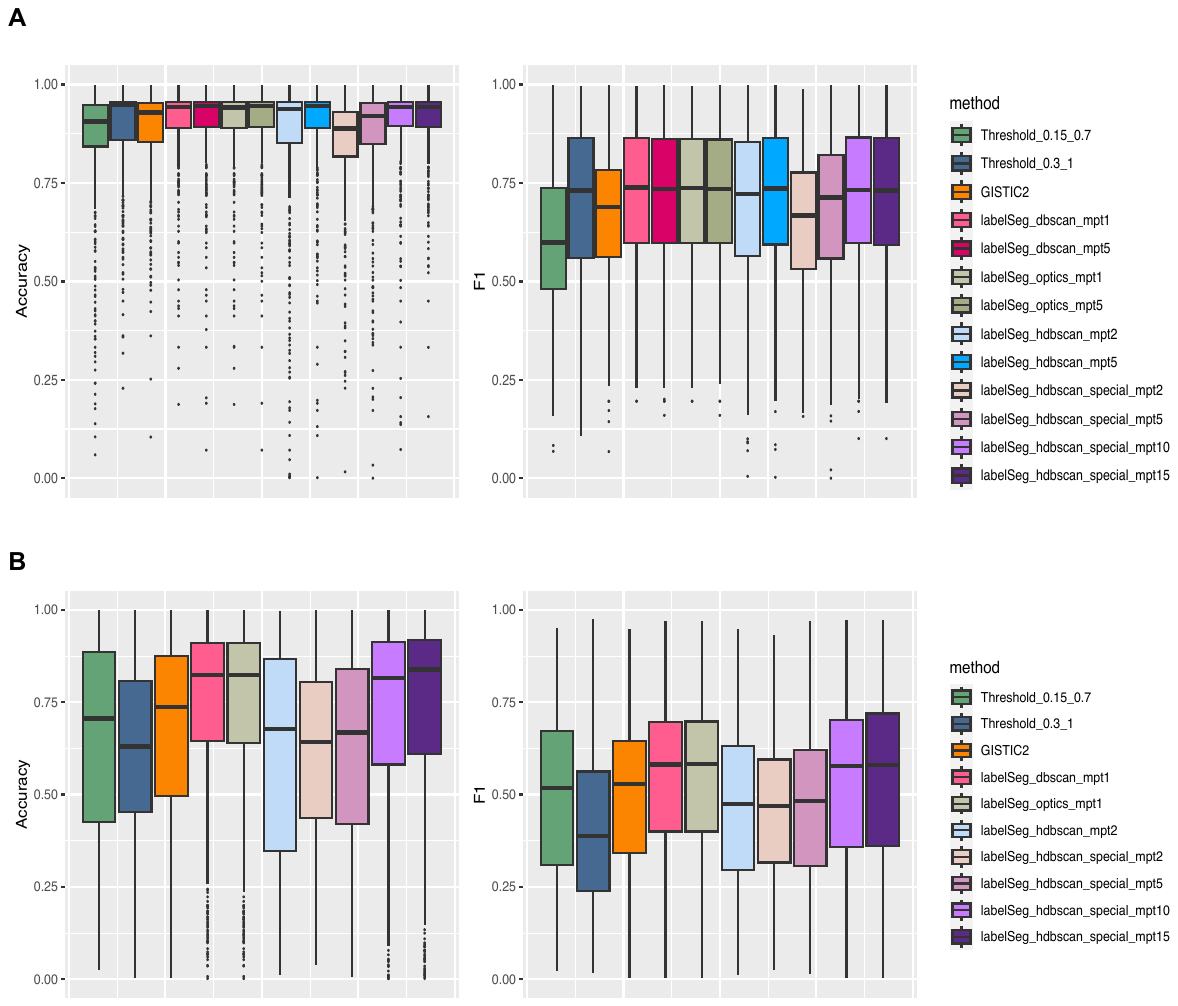

**Supplementary Figure S4: Benchmark performance on real data.** These plots illustrate the comparison between alternative methods and *labelSeg* with tuning in clustering methods and minPts parameters. For each sample, the macro average of F1 scores across all SCNA classes and overall accuracy were calculated. (A) Performance in the TCGA-GBM cohort. (B) Performance in the TCGA-LUSC cohort.

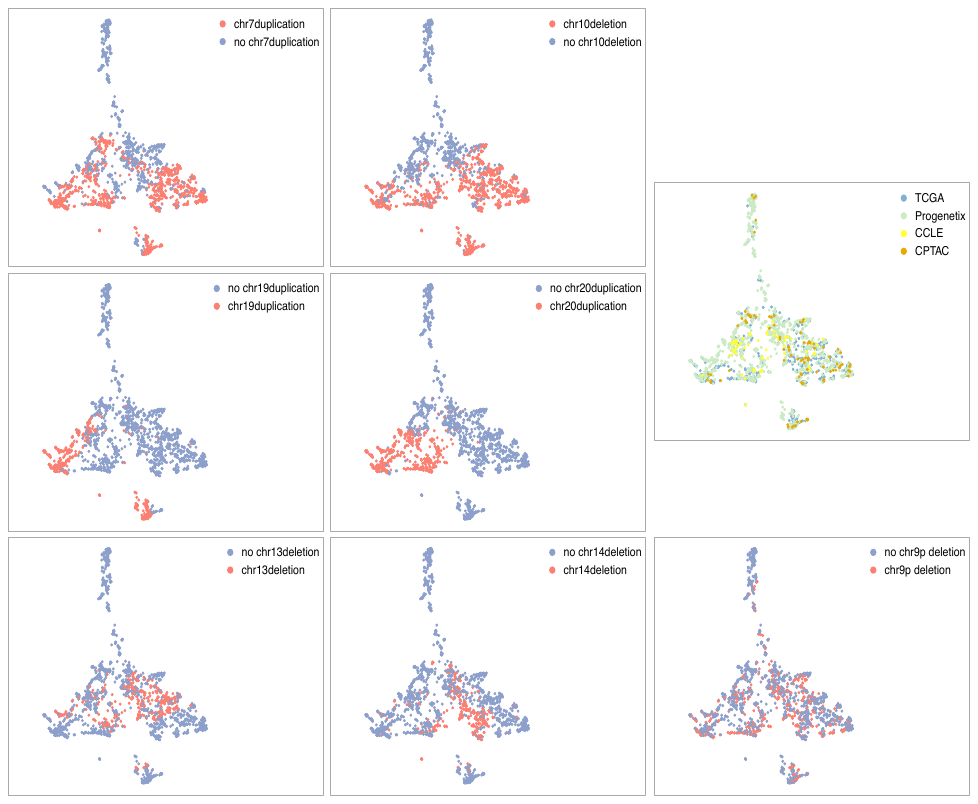

**Supplementary Figure S5:** **UMAP visualization of low-level SCNA coverage by chromosome for heterogeneous glioblastoma samples.** The plots are colored by chromosome SCNA (coverage by chromosome
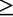
 70%), chromosome arm SCNA (coverage by chromosome arm > 50%), and data resources.

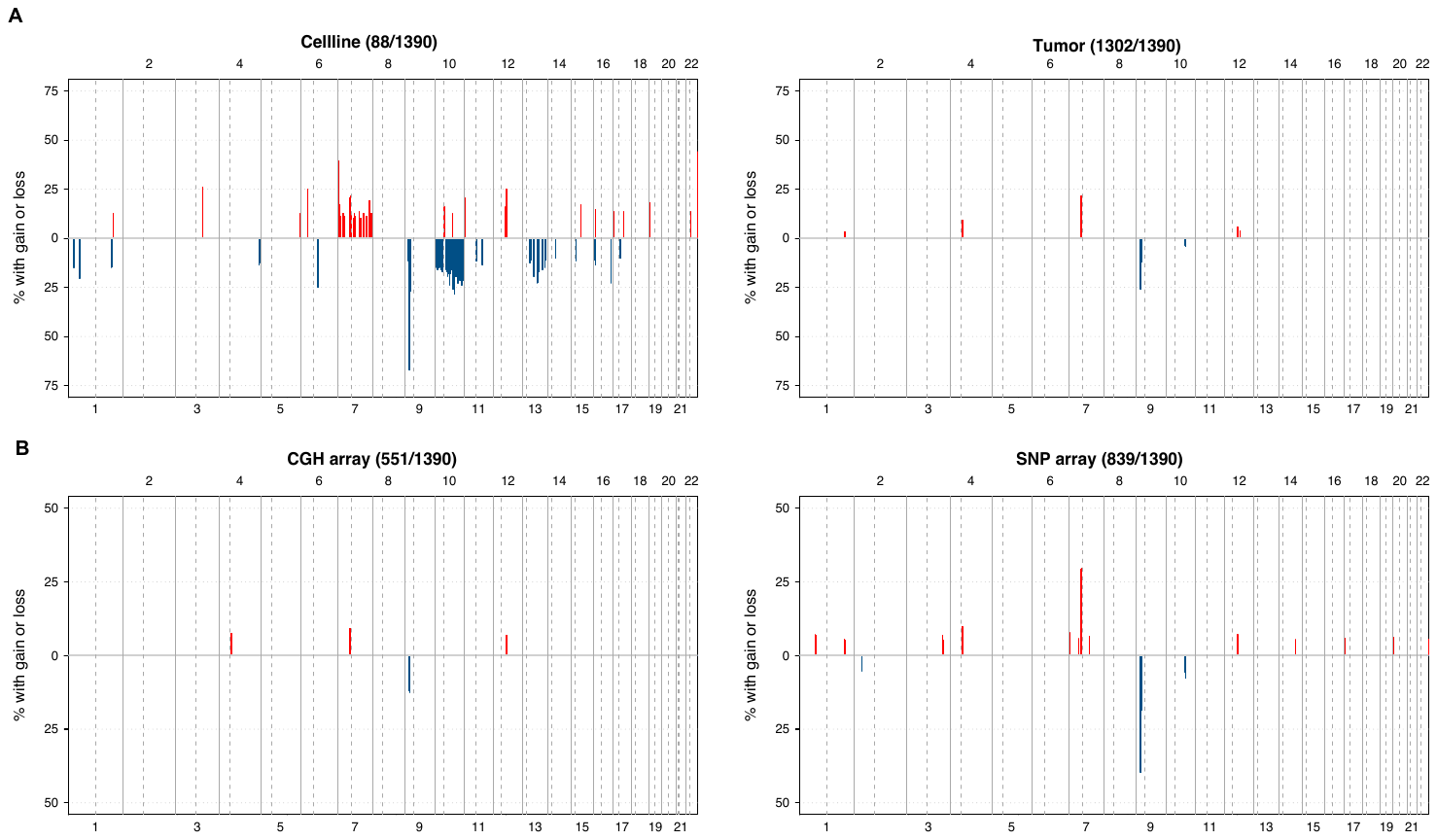
**Supplementary Figure S6: Frequency of high-level SCNA calls in the Progenetix dataset of glioblastoma samples.** (A) Analysis specific to sample types. (B) Analysis specific to array types. Background noise peaks were filtered out.

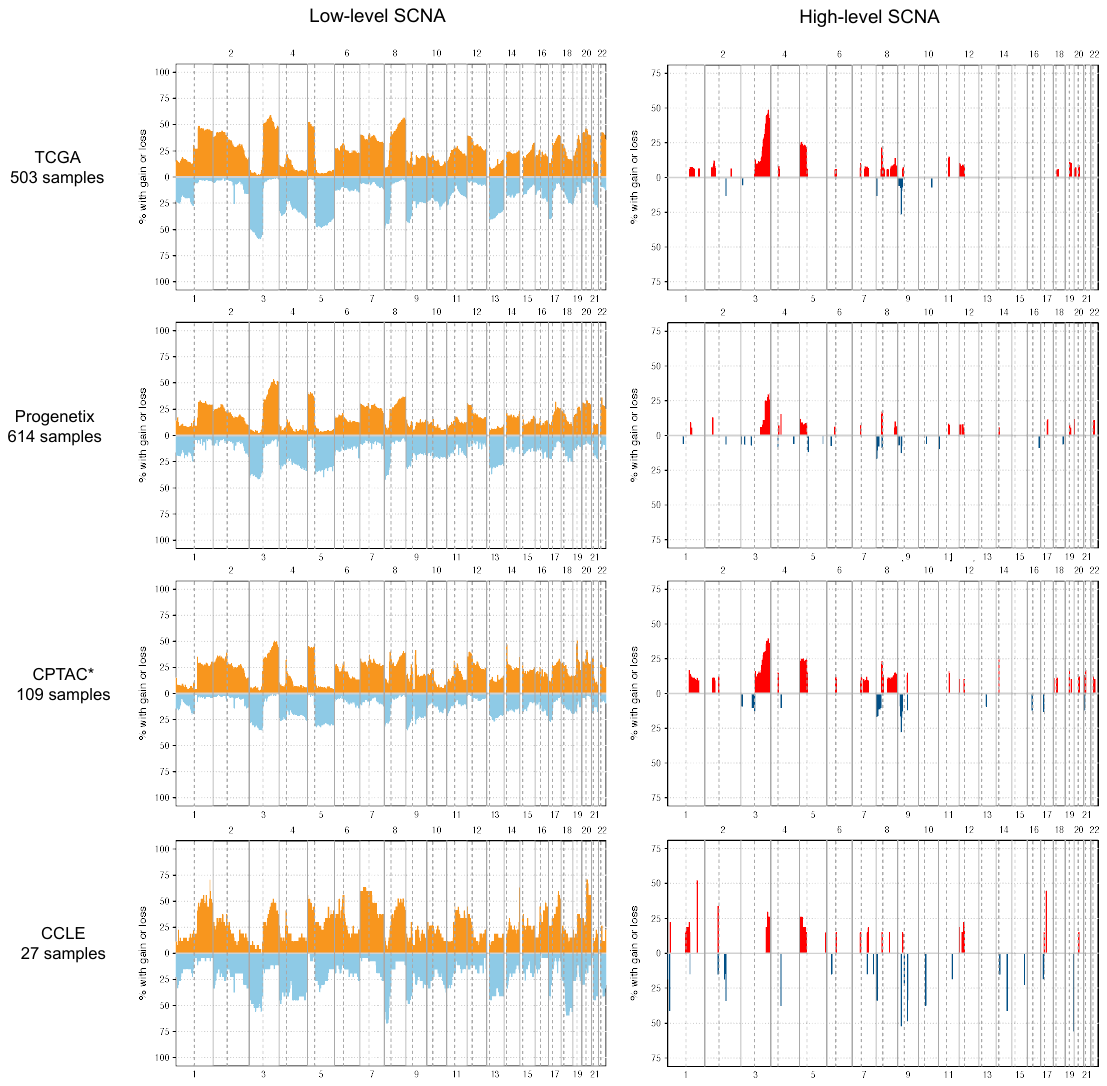

**Supplementary Figure S7: Frequency of SCNA calls in different datasets of lung squamous cell carcinoma samples.** Orange and red colors represent duplications, while light blue and dark blue colors represent deletions. The Y-axis displays the percentage of samples with SCNA overlapping with 1MB-sized genomic bins. The X-axis denotes chromosome numbers. Low-level SCNAs are segments identified by labels "+1" and "-1", while high-level SCNAs are segments denoted by labels "+2" and "-2". Background noise peaks were filtered out in the high-level SCNA frequency plots. It should be noted that, due to access limitations, the SCNA profiles of LUSC samples from CPTAC were obtained through ASCAT calls downloaded from the GDC data portal, rather than being generated by *labelSeg*.

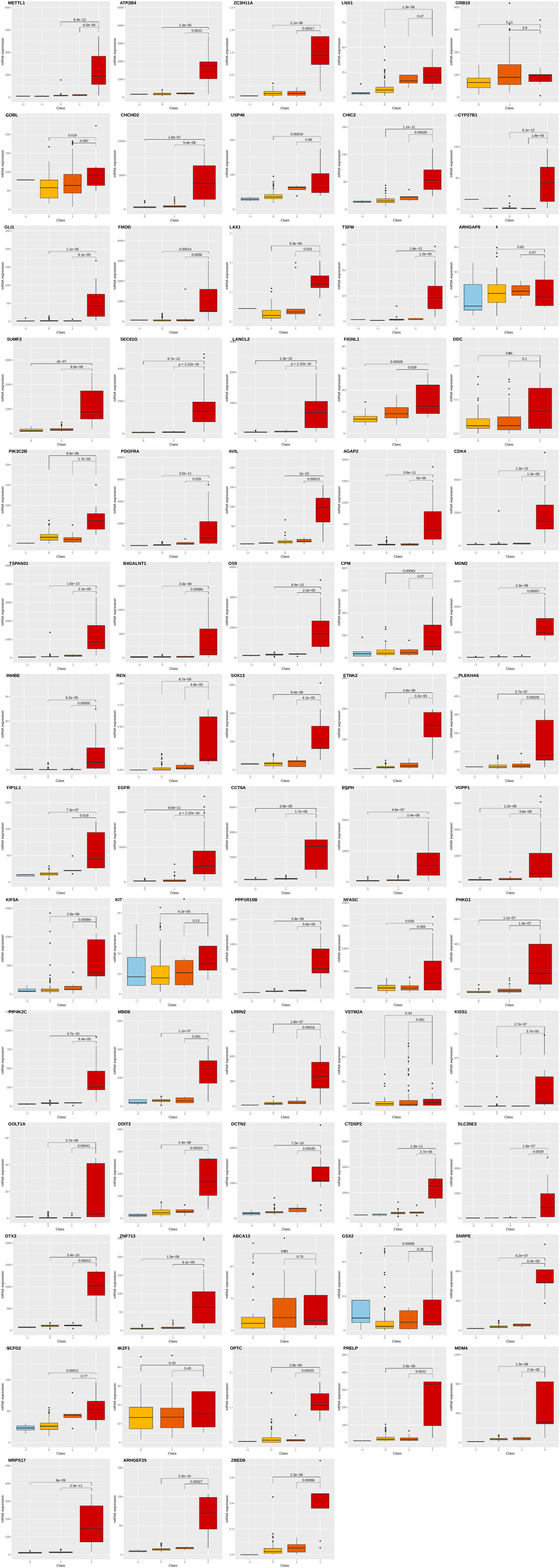

**Supplementary Figure S8: mRNA expression of genes with frequent high-level duplications in TCGA-GBM samples**

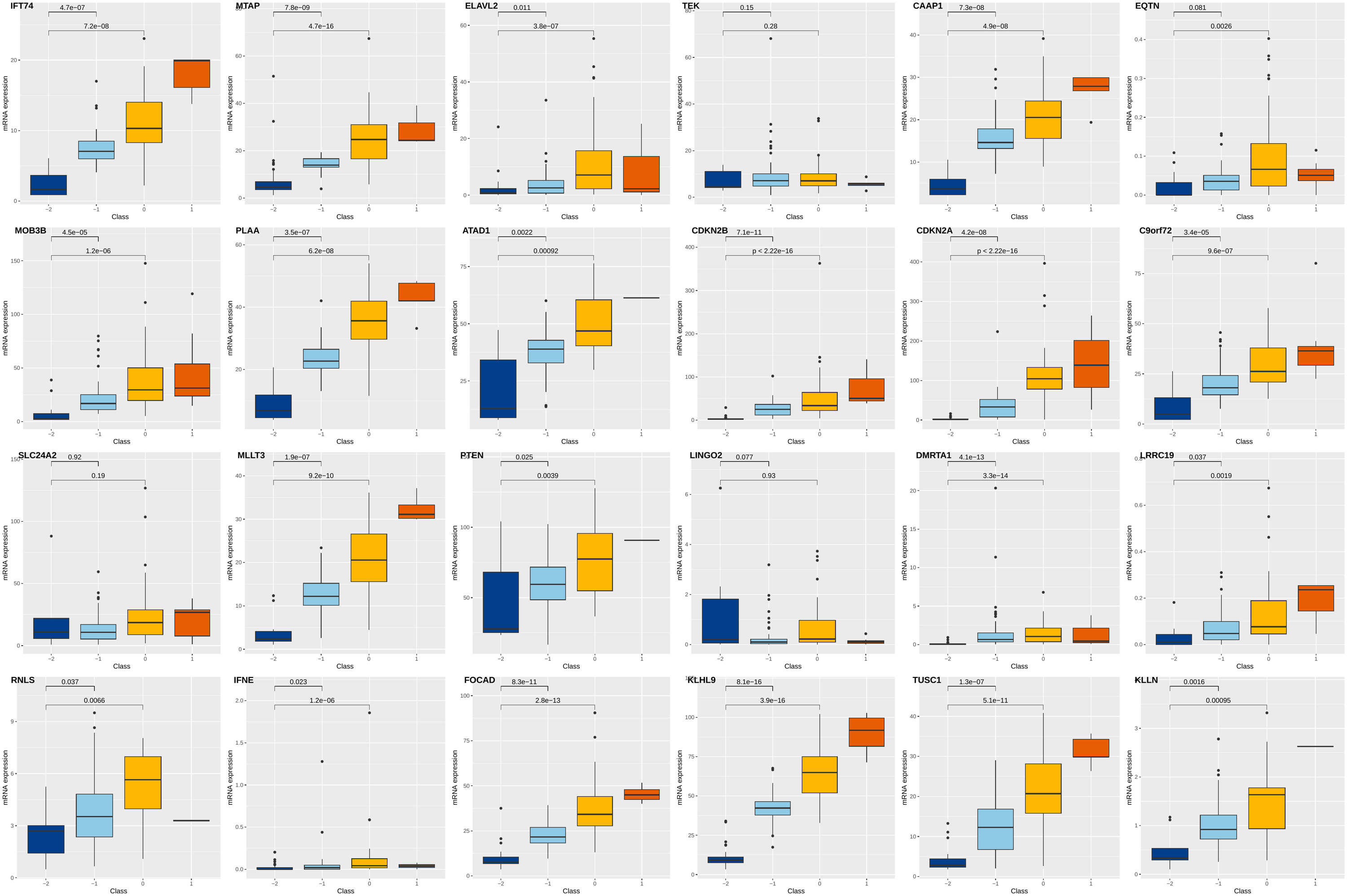

**Supplementary Figure S9: mRNA expression of genes with frequent high-level deletions in TCGA-GBM samples**

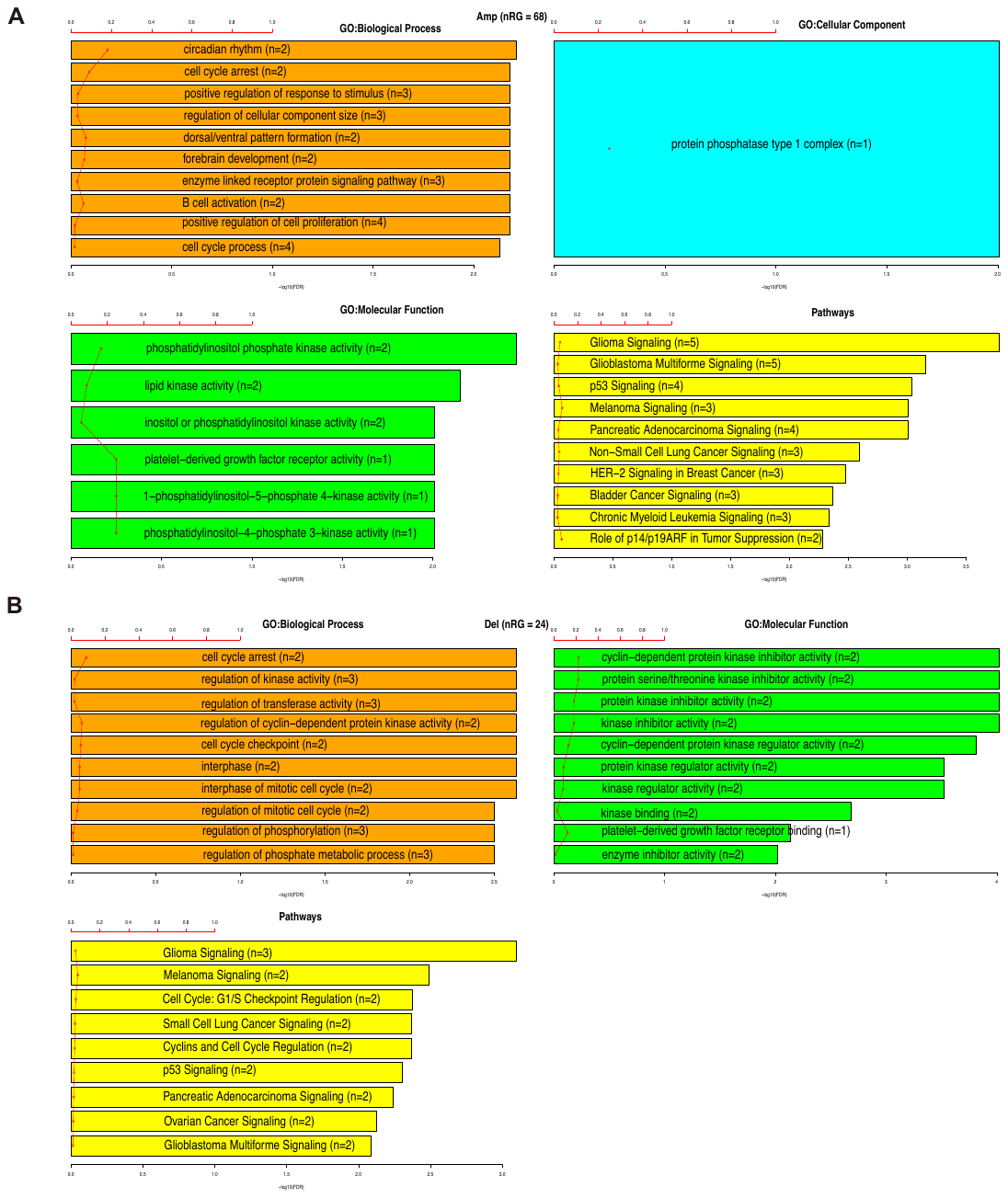

**Supplementary Figure S10: Enrichment analysis of genes frequently (A) high-level duplicated and (B) high-level deleted in TCGA-GBM samples.** Enrichment analyses were performed using the TCGAanalyze_EAcomplet function of the TCGAbiolinks R package.

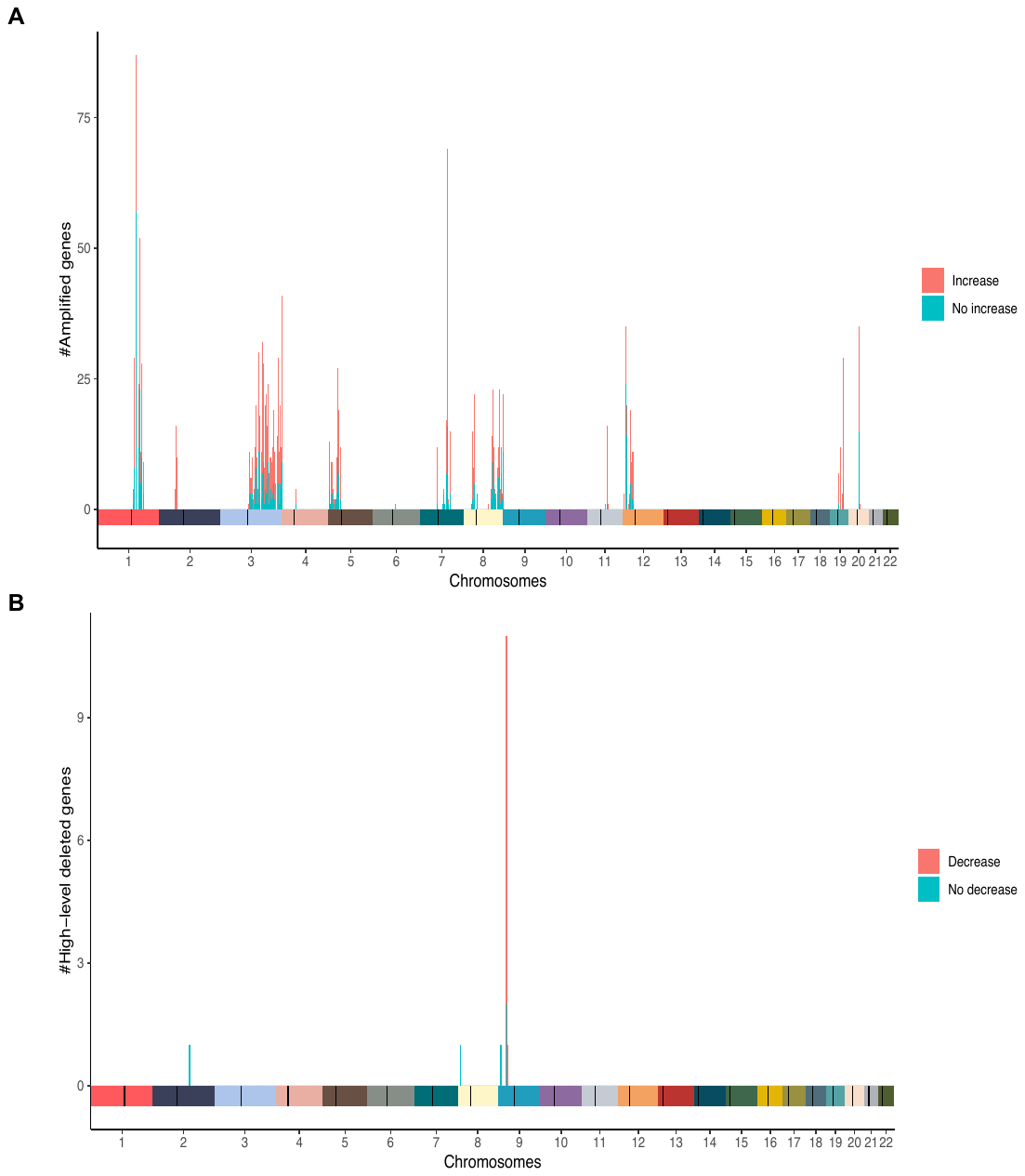

**Supplementary Figure S11: Genes frequently (A) high-level duplicated and (B) high-level deleted in TCGA-LUSC samples.** The protein-coding genes with an SCNA frequency greater than 5% in TCGA-LUSC samples and located in consensus high-level calling regions across at least two data resources were considered. The X-axis represents the genomic location indicated by chromosomes and centromeres. The Y-axis corresponds to the number of genes. Stacked bars are color-coded based on their association with mRNA expression, which was determined from matched TCGA-LUSC mRNA expression data.

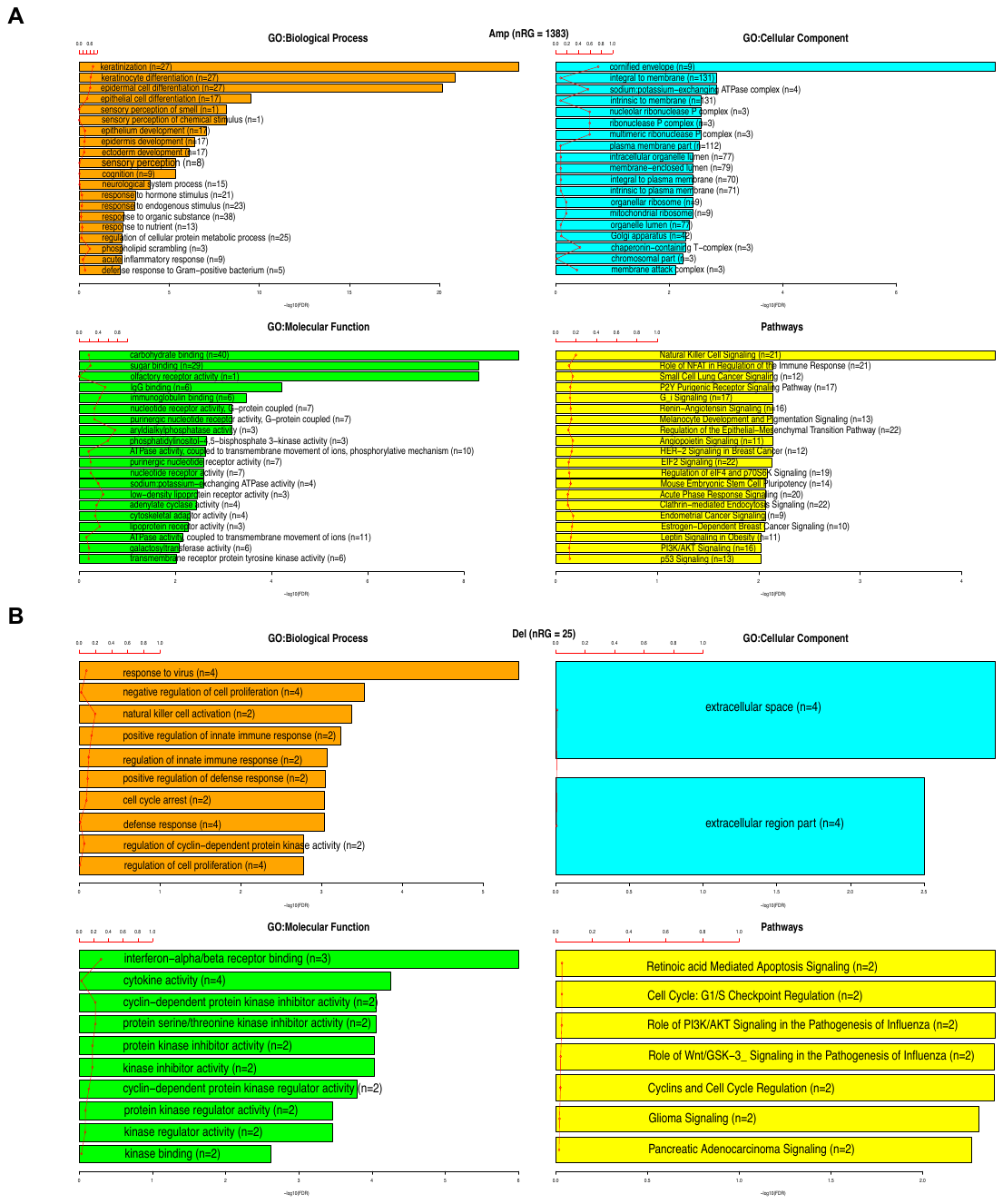

**Supplementary Figure S12: Enrichment analysis of genes frequently (A) high-level duplicated and (B) high-level deleted in TCGA-LUSC samples.**

### Reference

| [1] | H. Willenbrock and J. Fridlyand, "A comparison study: applying segmentation to array CGH data for downstream analyses," *Bioinformatics,* vol. 21, no. 22, p. 4084–4091, 2005. |
| --- | --- |
| [2] | A. Colaprico, T. C. Silva, C. Olsen, L. Garofano, C. Cava, D. Garolini, T. S. Sabedot, T. M. Malta, S. M. Pagnotta, I. Castiglioni, *et al.*, "TCGAbiolinks: an R/Bioconductor package for integrative analysis of TCGA data," *Nucleic acids research,* vol. 44, no. 8, pp. e71-e71, 2016. |
| [3] | E. Cerami, J. Gao, U. Dogrusoz, B. E. Gross, S. O. Sumer, B. A. Aksoy, Jacobsen, ers, C. J. Byrne, M. L. Heuer, E. Larsson, *et al.*, "The cBio cancer genomics portal: an open platform for exploring multidimensional cancer genomics data," *Cancer discovery,* vol. 2, no. 5, pp. 401-404, 2012. |
| [4] | J. Gao, B. A. Aksoy, U. Dogrusoz, G. Dresdner, B. Gross, S. O. Sumer, Y. Sun, Jacobsen, ers, R. Sinha, E. Larsson, *et al.*, "Integrative analysis of complex cancer genomics and clinical profiles using the cBioPortal," *Science signaling,* vol. 6, no. 269, pp. pl1-pl1, 2013. |
| [5] | C. Curtis, S. P. Shah, S.-F. Chin, G. Turashvili, O. M. Rueda, M. J. Dunning, D. Speed, Lynch, y. G, S. Samarajiwa, Y. Yuan, *et al.*, "The genomic and transcriptomic architecture of 2,000 breast tumours reveals novel subgroups," *Nature,* vol. 486, no. 7403, pp. 346-352, 2012. |
| [6] | K. M. Raine, P. Van Loo, D. C. Wedge, D. Jones, Menzies, rew, A. P. Butler, J. W. Teague, P. Tarpey, S. Nik-Zainal and P. J. Campbell, "ascatNgs: Identifying somatically acquired copy-number alterations from whole-genome sequencing data," *Current protocols in bioinformatics,* vol. 56, no. 1, pp. 15-9, 2016. |
| [7] | B. Gao, Q. Huang and M. Baudis, "segment_liftover: a Python tool to convert segments between genome assemblies," *F1000Research,* vol. 7, 2018. |
